# Dual role of E-cadherin in the regulation of invasive collective migration of mammary carcinoma cells

**DOI:** 10.1101/151613

**Authors:** Yair Elisha, Vyacheslav Kalchenko, Yuri Kuznetsov, Benjamin Geiger

## Abstract

In this article, we explore a non-canonical form of collective cell migration, displayed by the metastatic murine mammary carcinoma cell line 4T1. We show here that in sparsely plated 4T1 cells, E-cadherin levels are moderately reduced (~50%), leading to the development of collective migration, whereby cells translocate in loose clusters, interconnected by thin membrane tethers. Knocking down E-cadherin blocked tether formation in these cells, leading to enhancement of migration rate and, at the same time, to suppression of lung metastases formation *in vivo,* and inhibition of infiltration into fibroblast monolayers *ex vivo*. These findings suggest that the moderate E-cadherin levels, present in wild-type 4T1 cells, play a key role in promoting cancer invasion and metastasis.

## Introduction

Cancer invasion and metastasis is a highly versatile process, regulated at multiple levels, and characterized by several basic forms of cell migration ^1^. Recent studies carried out either *in vivo*, or in diverse *ex vivo* models, indicate that cancer cells utilize two major migratory strategies: keeping intercellular cohesion as collective or as single-cell invasion into the surrounding stroma ^2-6^.

The process of cancer cell individualization and acquisition of an invasive migratory phenotype commonly occurs within the framework of epithelial-mesenchymal transition (EMT), named after a similar process associated with morphogenetic transition during embryogenesis and organogenesis. During EMT, polarized epithelial cells undergo a major transcriptional and morphological transformation, resulting in the loss of their intercellular adhesions, and the acquisition of mesenchymal-like properties ^7–9^. As the EMT process progresses, the transformed cells lose the junctional connections with their neighbors, disengage from the epithelial layer in which they originated, and express mesenchymal markers such as N-cadherin, vimentin, and a multitude of specific transcription factors (e.g., Snail, Slug, Twist ^7, 10^). The acquired mesenchymal phenotype, is manifested by enhanced migratory activity, ECM production, invasiveness and elevated resistance to apoptosis ^7, 11, 12^. These changes enable the cells to penetrate into blood vessels and lymphatics and disseminate to distant organs, where they form cancer metastases ^1, 13^. Throughout this multi-stage process, the invading cells cross diverse tissue barriers such as basement membranes, fibrotic connective tissue, and vessel walls, using local degradation of the surrounding matrix by membrane-bound or secreted proteases ^1^ and development of actin-based invadopodia ^14, 15^, which combine mechanical and enzymatic activities to penetrate into the pericellular matrix ^11^.

Microscopy-based live-cell imaging *in vivo* and diverse *ex vivo* models have demonstrated that loss of tissue integrity is usually followed by two major forms of single-cell migration: an amoeboid migration, most prominently displayed by highly anaplastic tumors; and mesenchymal migration, commonly displayed by sarcomas, glioblastomas and carcinomas ^16–18^. It was further shown that tumor cells can switch between these two modes of migration ^19, 20^.

Cells that demonstrate amoeboid movement are characterized by relatively high migratory speed ^17, 18, 21^, poor adhesion to the extracellular matrix (ECM) ^17, 18, 21^, and a high degree of deformability, which, collectively enable them to cross physical tissue barriers. Invasive mesenchymal migration, on the other hand, is relatively slow ^3, 6, 17^, requires adhesive interactions with the ECM, usually via integrin receptors ^3, 6^, and commonly uses matrix metalloproteinases to destroy local ECM barriers encountered during the migration ^3, 6^.

It is noteworthy that EMT can be an intrinsic, “cell autonomous” process, caused by genetic alterations taking place within the primary tumor ^13, 22^, or an environmentally-driven process, induced by stromal or inflammatory cells located within or around the primary tumor ^13, 22^. The former mechanism is at work mainly in primary, undifferentiated carcinomas, and thought to be associated with a stable, “EMT-stemness” state ^22^, whereas environmentally-induced EMT is often a transient process, enabling the cells to re-acquire epithelial properties following mesenchymal-epithelial transition (MET) ^9, 12, 22, 23^.

Collective migration is another hallmark of invasive epithelial cancers, characterized by the capacity of cancer cell assemblies, interconnected by stable cell-cell junctions (mostly cadherin-mediated adherens-type junctions), to move through the ECM together, while maintaining their cell-cell connections ^2–4, 6^. The migration of cells as a coherent entity requires a high level of intercellular coordination, a process attributed to cytoskeleton-mediated mechanical coupling between neighboring cells ^2, 24, 25^. The migrating cell collectives display clear anterior-posterior polarity, whereby a group of cells serves as the “invasive front”, while those at the opposite end follow them ^2-4, 25, 26^.

The mechanisms regulating collective migration, and their roles in metastasis, are still poorly understood. Notably, collective migration commonly leads to local invasion of cancer cells, while the formation of distant metastatic lesions requires loss of intercellular coherence, enabling individual cancer cells or small cell clusters to dislodge from the primary tumor, penetrate into blood vessels or lymphatics, and extravasate into distant organs, forming a metastatic lesion ^1, 9, 12, 13, 27^.

In this study, we explored a “non-canonical” mechanism of collective cell migration displayed by the highly metastatic murine mammary gland carcinoma cell line, 4T1 cells. When injected either into the mammary fat pad or into the bloodstream, these cells develop metastases in the lung, liver, and bone, and serve as an animal model of highly metastatic breast cancer ^28, 29^. Interestingly, despite their invasive phenotype, 4T1 cells in confluent cultures express high levels of E-cadherin ^30–32^.

We further show here that in sparsely-growing 4T1 cells, E-cadherin levels are post-transcriptionally down-regulated and the cells form loose clusters, interconnected by long, thin membrane protrusions (“tethers”). Live-cell imaging demonstrated that the cells within each cluster have considerable freedom to migrate individually; yet their departure from the cluster is restricted by the tether. Examination of lung metastases formed from 4T1 cells, using two-photon microscopy, revealed multiple tethers interconnecting the tumor cells. Furthermore, we found that each tether extends from a single cell, which is attached to neighboring cells via Ecadherin-rich adhesions. Knockdown of E-cadherin blocked tether formation, and switched the motility mode of the cells from collective to single-cell migration. We show here that *ex vivo,* E-cadherin mediates tether formation, and facilitates 4T1 cell infiltration into the stroma; while its knockdown significantly reduces the metastatic dissemination of 4T1 cells to the lungs, following intravenous or fat-pad injection.

## Results

### Characterization of tether-mediated collective migration of cultured 4T1 cells

In a search for cellular mechanisms underlying invasive cell migration, we examined various epithelial cancer cell lines for their migratory characteristics, using live-cell imaging. Sparsely plating these cells (to reduce the frequency of random cell-cell interactions), we examined the tendency of the different cell lines tested to migrate collectively, by calculating the average “nearest neighbor” distances between them (see examples in Supplementary Fig. S1 and Supplementary Movie S1). Some of the cell lines (e.g., A431 and MDCK) displayed poor migratory activity in the absence of external stimulation, and had a tendency to form rather coherent and stationary islands, while MCF10A cells formed small epithelial islands, displaying slow but persistent migratory activity (Supplementary Movie S1). Among the other cell types tested, the highest levels of persistent migratory activity were seen in H1299 cells, which primarily migrated as single cells, and exhibited a broad range of largely dispersed cell-to-cell spacings (on the order of 50-240 μm; Supplementary Fig. S1B), and 4T1 cells, which displayed a unique form of collective cell migration (cell-to-cell spacings of around 20-30 μm; Supplementary Fig. S1B; Supplementary Movie S1).

Time-lapse imaging of co-migrating 4T1 cells indicated that the cells do not display extensive cell-cell junctions, similar to those of MDCK and MCF10A cells, but are rather interconnected via an elaborate network of thin membrane tethers (Fig. 1A; Supplementary Movie S1). These co-migrating clusters of 4T1 cells are formed partly by cell-cell interactions following cell division, and partly by merger with neighboring cells or clusters (Supplementary Movie S2). Based on scanning electron microscopy (SEM) and light microscopy morphometry, the length of the tethers varies from around 20 μm to more than 100 μm, while the width of their shafts varies between 20 and 200 nm.

**Figure 1:**
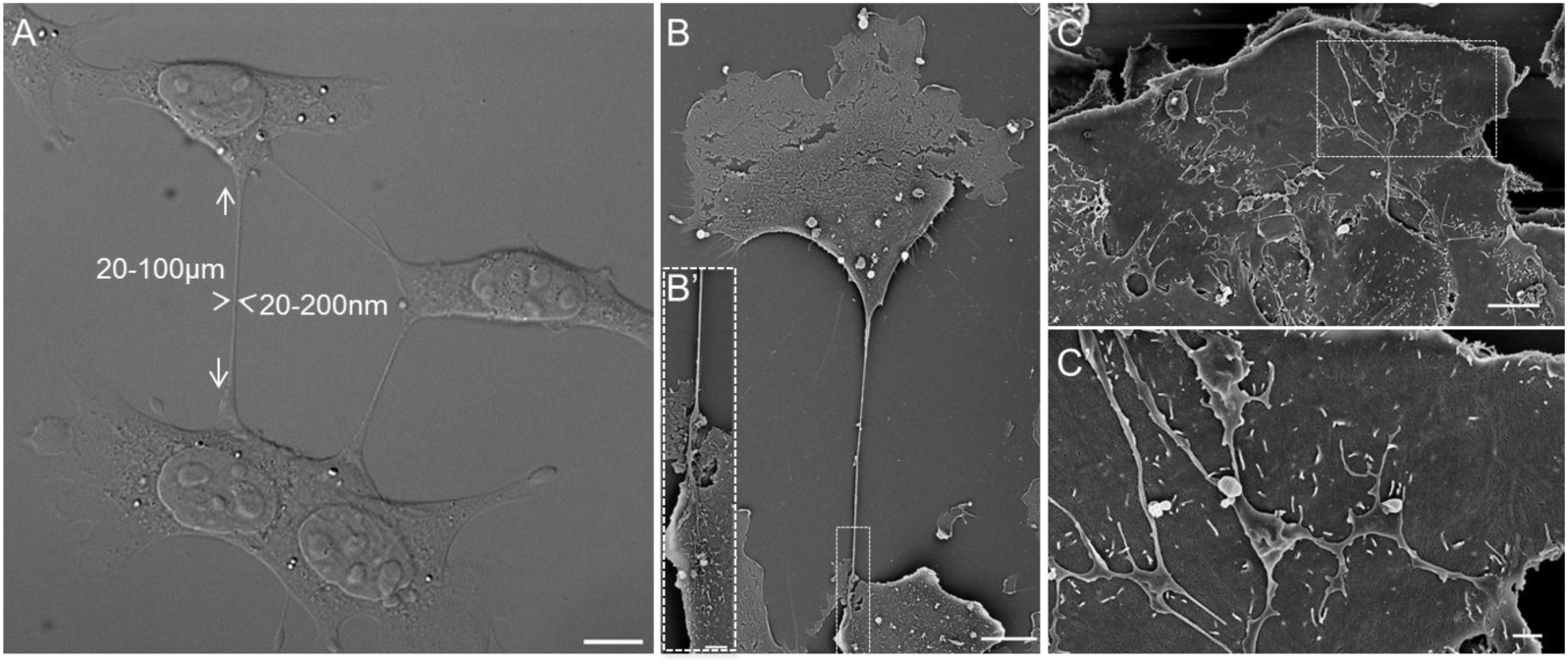
Formation of intercellular tethers in sparsely and densely plated cell cultures. 4T1 cells were sparsely (A and B) or densely plated (C and D) on tissue culture dishes, coated with 10 μg/ml fibronectin (FN, and incubated for 24 hrs. The attached cells were then fixed and observed, using Nomarsky optics (DIC) (A) or scanning electron microscopy (SEM) (B). The images show multiple, membrane tethers that interconnect neighboring cells. Typical dimensions of the tethers are indicated by arrows and arrowheads in A. Inserts B’ and C’ display the marked regions in B and C, respectively, at higher magnification. It is noteworthy that in sparse cultures, cells display 1-2 unbranched tethers; in dense cultures, multiple branched tethers are seen. Scale bars: A and B=10 μm; B’=2 μm; C=5 μm; C’=1 μm.

Moreover, live-cell imaging demonstrated that the tethers are asymmetrical, formed when a cell begins to pull away from a neighboring cell, while still maintaining limited intercellular adhesion that keeps the two cells tether-connected (Supplementary Movies S1 and S3). Differential interference contrast (DIC) microscopy (Fig. 1A) and SEM examination (Fig. 1B and B’) demonstrated that in sparse cultures, 4T1 cells tend to form a small number of tethers (usually 1-3), while in dense cultures, extended adherens junctions are formed, accompanied by a network of branched membranal protrusions that bridge between neighboring cells (Fig. 1C and C’). End-to-end examination of individual tethers indicated that each tether originates in one cell and is attached, via its opposite end, to the other cell’s membrane (Fig. 1B and B’).

To explore the functionality of the tethers as regulators of collective migration, we utilized live-cell video microscopy, which enabled us to monitor the formation and fate of individual tethers. An unbiased assessment, based on the monitoring of over 100 tethers, indicated that about 50% of them completed a “full departure-return cycle”. A similar analysis carried out with H1299 cells indicated that the migrating cells produced highly unstable tethers, which were maintained only in about 4% of the cases. To determine the effect of tether-mediated cell-cell interactions on migration speed, we subjected movies of migratory 4T1 cells to quantitative analysis, and compared the migration rates of clusters containing different cell numbers. The results, shown in Fig. S2, clearly indicate that small cell cluster ( containing 2-7 cells), translocate ~1.6-fold faster than larger clusters (consisting of over 8 cells), and, interestingly, also 3.7-fold faster than single 4T1 cells.

Measurement of the lengths of the different tethers produced by those 4T1 cells that eventually returned to their sites of origin is shown in Fig. 2A. In most cases, the rates of tether extension and shortening were similar, and the dwell time between departure and return was roughly proportional to the maximal tether length. Tracking of individual tethered cells over time (Fig. 2B; Supplementary Movie S3) revealed that departing cells develop a lamellipodium at the edge of the cell, opposite the tether, but after reaching its maximal “outward migration” (Stage 4 in Fig. 2A and B), the position of the lamellipodium is often reversed, and a new one develops in the vicinity of the tether, guiding the cell back to its original location (Stages 6-9 in Fig. 2).

**Figure 2:**
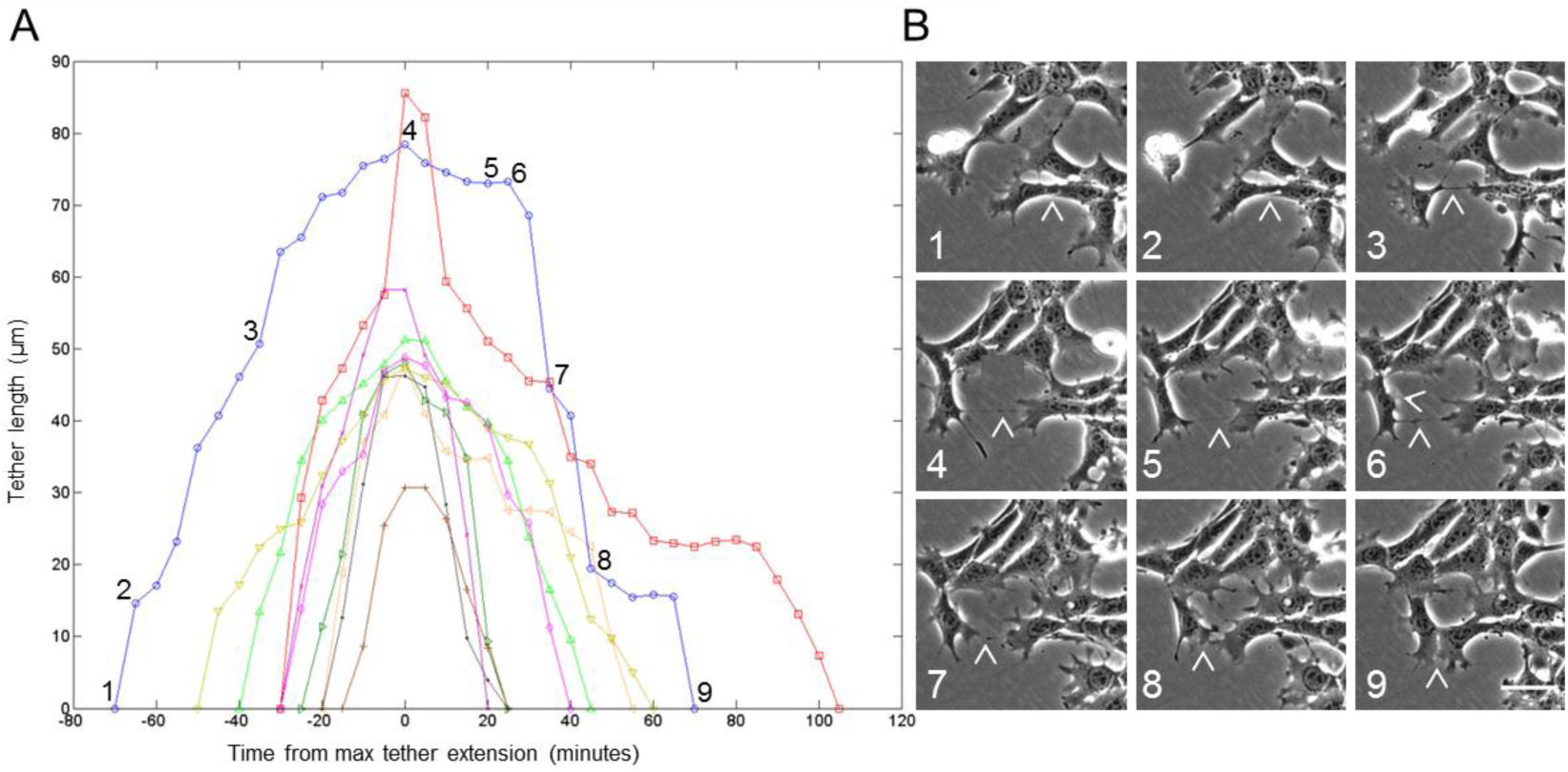
Dynamic modulation of tether formation and length. (A) 4T1 cells were cultured for 24 hrs on tissue culture dishes coated with 10 μg/ml fibronectin (FN). Cell movement was then monitored by live-cell phase contrast microscopy. Using these movies, we measured the formation, elongation, and shortening of individual tethers, over time. Each curve represents changes displayed by a single tether. The blue curve, and the associated numbers, correspond to the temporal snapshots shown in (B). Arrowheads denote the measured tether, the features of which are described in Results. Scale bar: 50 μm.

### Cytoskeletal and adhesive properties of 4T1 intercellular tethers

The results presented in Figure 2 raise the possibility that the interconnecting tethers mechanically restrain the outward movement of cells within each cluster, a property that might require considerable tensile strength on the part of the tether, largely determined by its intrinsic resistance to rupture, and the strength of its adhesion to the neighboring cell.

To explore the structural properties of the tether, we examined their cytoskeletal contents by means of immunofluorescence microscopy. The results, shown in Figure 3, indicate that the tethers contain all three cytoskeletal components; namely, F-actin, microtubules, and vimentin-containing intermediate filaments (Fig. 3A, B and C). Labeling of densely-plated 4T1 cells for E-cadherin and β-catenin revealed circumferential adherens junctions, as well as multiple patches along the dorsal cell membrane, containing the two proteins (Fig. 4A,C). These dorsal patches most likely correspond to the adhesions of the branched protrusions (Fig. 1C and C’). Sparsely-plated 4T1 cells displayed reduced E-cadherin levels, mostly located at the tether attachment sites (Fig. 4B). We further found that the β-catenin-rich adhesions at the tethers’ ends also contain prominent levels of actin (Fig. 4D).

**Figure 3:**
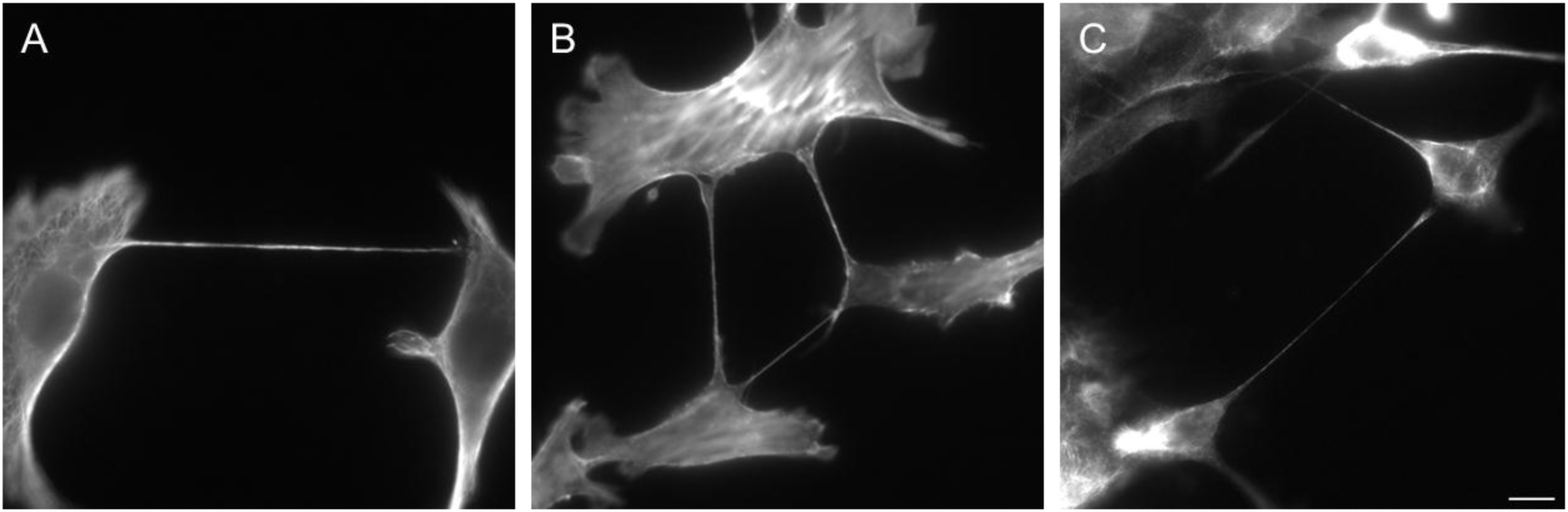
Cytoskeletal support of 4T1 cell tethers. 4T1 cells were cultured for 24 hrs in tissue culture dishes, then fixed and immunolabeled for Tubulin (A), Actin (B), or Vimentin (C). Scale bar: 10 μm.

β-catenin and cadherin are hallmarks of calcium-dependent adherens-type junctions and, indeed, addition of the calcium chelator EGTA to the culture medium induced tether retraction (Supplementary Fig. S3; Supplementary Movie S4). To quantify the effect of Ca^++^ chelation and reconstitution on tether formation, we determined the number of tethers per cell at different time points following the addition of EGTA to the medium, and after Ca^++^ restoration. As shown in Fig. S3, EGTA treatment initially induced cell contraction, leading to the formation of multiple short-lived tethers (t=5 min), which essentially disappeared upon further incubation (t=60 min). Following restoration of normal Ca^++^ levels, a gradual increase in tether number was measured, reaching normal levels by 60-120 min of incubation.

Comparison of E-cadherin levels in densely- and sparsely- plated 4T1 cells using Western blot analysis revealed a twofold lower level of E-cadherin in the sparsely plated cells (Fig. 4E and E’). Average E-cadherin RNA levels, on the other hand, were not affected by culture density (Fig. 4F), indicating that the density dependence of E-cadherin levels is regulated post-transcriptionally. To further characterize the effect of cell density on E-cadherin levels, we plated 4T1 cells in decreasing numbers on 2D surfaces, and quantified their E-cadherin content by Western blot analysis. As shown in Fig. S4, the level of E-cadherin reaches its maximal value at nearly 100% confluence, and gradually decreases at lower cell confluence levels. At ~10% confluence or lower, the levels of E-cadherin displayed decline twofold, compared to those of confluent cells. These reduced levels are comparable to those shown in Figure 4E and E’.

**Figure 4:**
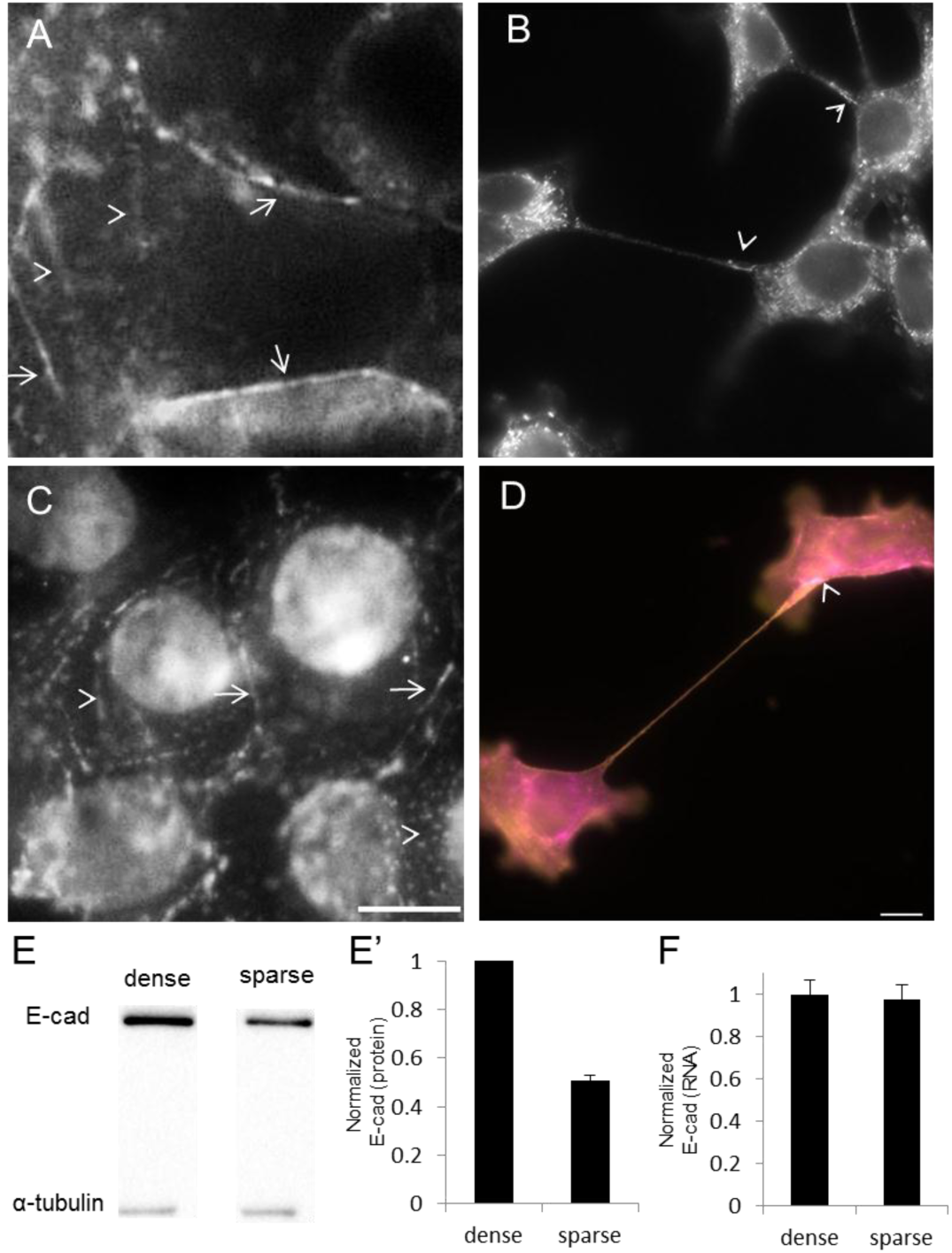
Tether-mediated cell-cell adhesions are mediated via adherens-type junctions. (A,C) Densely-plated 4T1 cells form E-cadherin- (A) and β-catenin-rich (C) adherens junctions, located either along the sub-apical region (arrows), or in multiple dorsal patches (arrowheads). (B,D) Sparsely plated 4T1 cells form structurally and molecularly polarized tethers that are attached to the neighboring cell via E-cadherin (B), and co-localized actin (magenta) and βcatenin (yellow) (D). Arrowhead in panel D denotes the tether adhesion area. Scale bar: 10 μm. (E, E’) Immunoblotting and corresponding histogram depicting E-cadherin levels in sparsely-(5.6*10^3^/cm^2^) and densely-Plated (8.9*10^4^/cm^2^) 4T1 cells. The normalized levels of E-cadherin (relative to α-tubulin) are shown in E’. (F) Comparison of E-cadherin mRNA levels in densely- and sparsely-plated 4T1 cells normalized to HPRT1 and GUSB genes. The results are based on two independent experiments.

To test whether tether formation can be induced in cells that lack E-cadherin, by moderate induced expression of this adhesion molecule, we transfected H1299 cells stably expressing a tetracycline-inducible E-cadherin vector (a kind gift from Barry M. Gumbinar, Seattle Children’s Research Institute, Seattle, Washington, USA) with the transfected H1299 cells. We show that induced expression of E-cadherin at levels comparable to those expressed in sparse 4T1 cells, resulted in the formation of stable tethers in H1299 cells, leading to the acquisition of collective migratory activity (Supplementary Fig. S5 and Supplementary Movie S5). A complementary approach involving treatment with moderate concentrations of EGTA indicated that MCF10A cells, which normally exhibit “classical”, slow collective migration, lost their tight cell-cell adhesion and migrated collectively, as tether-inter-connected clusters, following such treatment (Supplementary Fig. S6; Supplementary Movie S6).

To directly assess the role of E-cadherin in tether formation, 4T1 cells were subjected to Ecadherin knockdown (shEcad), reducing E-cadherin mRNA levels by 85%, compared to non-targeting control shRNA (shCon), and inducing a threefold reduction in tether formation (Fig. 5 and Supplementary Movie S7). It is noteworthy that loss of tethers upon E-cadherin knockdown was also noted in 4T1 cells growing in 3D collagen gels (Supplementary Movie S8).

**Figure 5:**
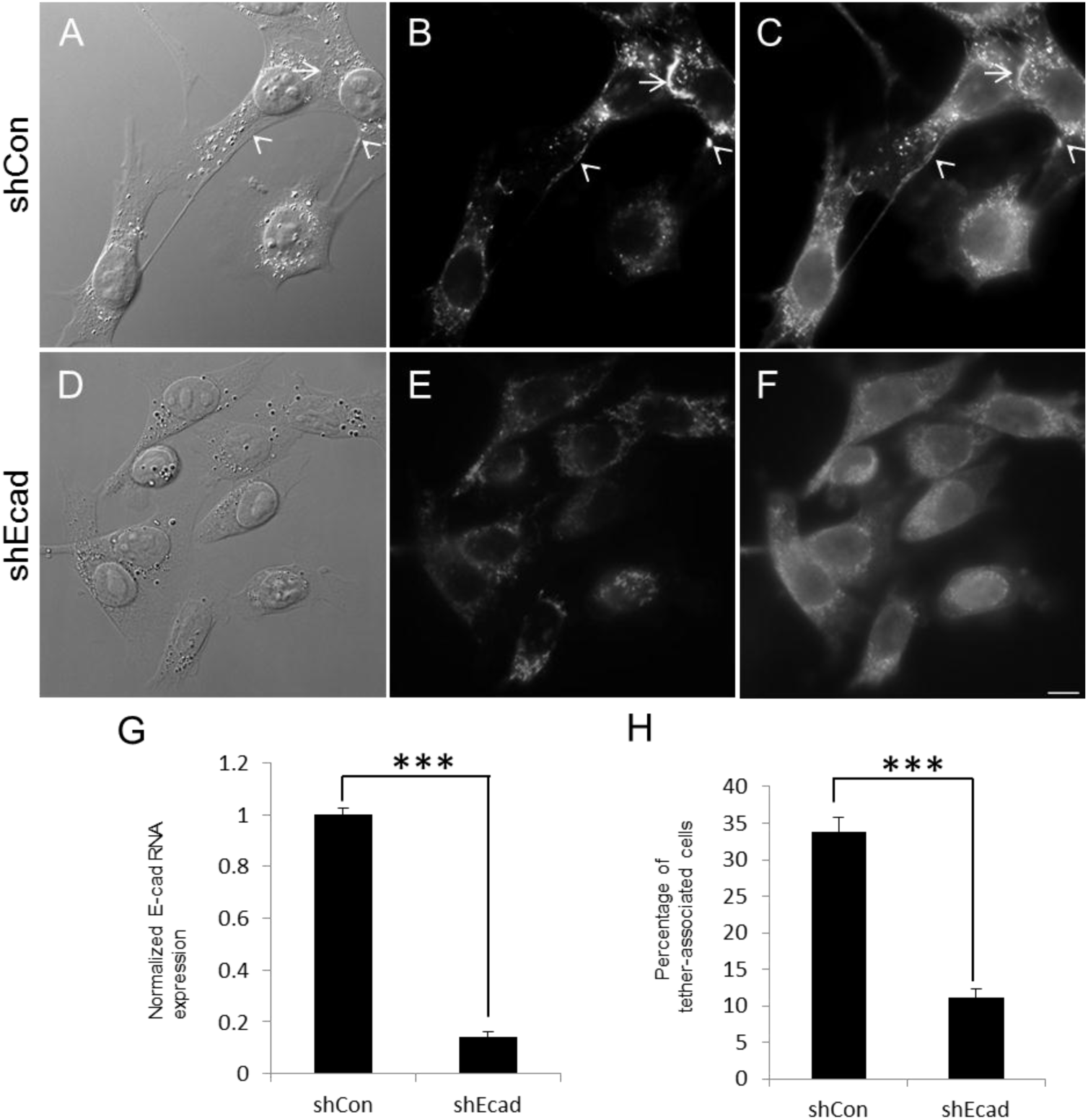
E-cadherin is essential for tether formation. 4T1 cells, stably knocked down for E-cadherin (shEcad), and control cells (ShCon) were imaged by DIC (A, D) and double-labelled for E-cadherin (B, E) and β-catenin (C, F). Note that tether formation and E-cadherin-rich adhesions are markedly reduced in the knocked-down cells. (G) qRT–PCR analysis, pointing to an over 85% reduction in E-cadherin mRNA levels in shEcad compared to shCon 4T1cells, obtained in 3 independent experiments. (H) Comparison of the percentage of tether-associated cells in shEcad and shCon 4T1cells, pointing to ~68% reduction in the knocked-down cells, compared to controls. ***p < 0.001. Scale bar: 10 μm. n=11 different images (containing around 800-900 cells in total for each group), obtained in two independent experiments.

### The involvement of E-cadherin in the formation of 4T1 lung metastases

Sparsely plated 4T1 cells manifest multiple cellular and molecular features that are hallmarks of the epithelial-mesenchymal transition (EMT), including the loss of both apical-basal polarity and adherens junctions (Figs. 1B, B’ and 4B), as well as the expression of vimentin, and acquisition of a migratory phenotype ^7, 9, 12^. These observations are in line with the notion that reduced expression of E-cadherin in these particular cells leads to the loss of coherent adherens junction formation; yet its levels remain sufficient for supporting tether-mediated collective migration (see Discussion, below).

To explore the involvement of the residual E-cadherin present in 4T1 cells on their metastatic properties, *in vivo*, we expressed either green fluorescent protein (GFP), for 2-photon microscopy, or luciferase, for IVIS bioluminescent whole body monitoring of these cells in BALB/c mice. The cells were injected into the tail vein or the mammary fat pads of the mice. Preliminary examination of lung metastases, formed 7 days after tail vein injection of 4T1-GFP-shCon cells, revealed loosely packed metastatic nodules containing cells interconnected by multiple tethers (Fig. 6A; Supplementary Movie S9), comparable to those found in 4T1 cells, cultured in 3D collagen gels (Supplementary Movie S8). These observations confirm that tether formation occurs *in vivo*, and is compatible with formation of metastases.

**Figure 6:**
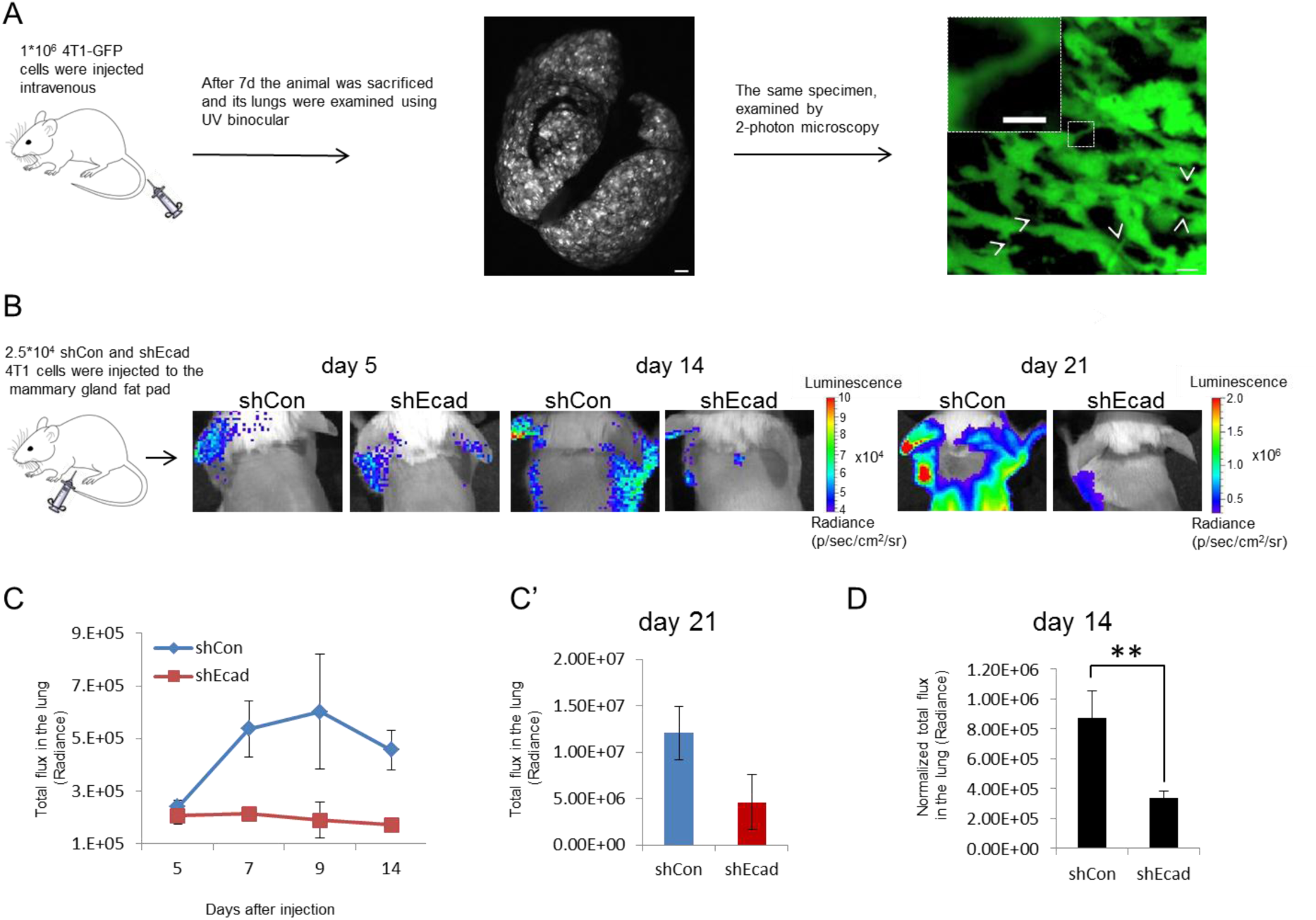
Effect of E-cadherin Knockdown on the formation of lung metastases following injection of 4T1 cells into BALB/c mice. (A) 4T1-GFP-shCon cells (1*10^6^ cells) were injected intravenously into the tail veins of BALB/c mice. Seven days later, animals were sacrificed, and their lungs imaged using 2-photon microscopy. Examination of the images reveals loosely packed metastatic nodules with multiple tethers running between the cells (arrowheads). Scale bar: Middle image - 10 mm; right image - 10 μm; insert - 5μm.(B) Representative bioluminescence images (BLI), taken on days 5, 14, and 21, post Injection of 2.5 *10^4^ 4T1-luciferase shCon or shEcad cells into the mammary fat pads of BALB/c mice (n=4/5 mice, in each group, respectively). (C, C’) BLI quantification of total flux in the lungs of shCon or shEcad mouse groups. Note that the accumulation of tumor cells in the lungs during the 14/21 day post-injection was over twofold lower in the E-cadherin knocked-down cells, compared to the control 4T1 cells. (D) BLI quantification of total flux of 4T1-luciferase shCon or shEcad, in the lungs of BALB/c mice studied in two separate fat-pad injection experiments (n=9/10 for each shCon or shEcad group, respectively). Data were collected on Day 14, and statistically significance of the differences between the groups was determined, using two-way ANOVA (**p= 0.009).

Interestingly, comparison of the metastatic activity of control 4T1 cells (shCon) with that of E-cadherin knockdown cells (shEcad), indicated that suppression of E-cadherin expression markedly reduced the metastatic dissemination of the cells. Specifically, luciferase-expressing 4T1-shCon cells or E-cadherin knockdown cells (4T1-shEcad) were injected, separately, into the mammary fat pads of BALB/c mice (Fig. 6B), and the formation of lung metastases was quantified by a non-invasive bioluminescence assay. As shown in Figure 6B and C, both groups formed metastatic lesions; yet the bioluminescence in the lungs, measured on Day 7 post-injection, was ~2.5-fold higher in 4T1-shCon mice, compared to the 4T1-shEcad group. This difference remained consistent over time, up to 2 weeks post-injection. During the third week post-injection, a major increase in lung-associated bioluminescence was noted in both the shCon and shEcad mice groups, which may be attributed to massive cancer cell proliferation in the lungs. Notably, also at that stage, the ~2.5-fold difference between the two groups remained unchanged (Fig. 6C’).

To determine the significance of the effect of E-cadherin knockdown on 4T1 metastasis, the bioluminescence results were subjected to two-way ANOVA, which indicated that the difference between the two groups is highly significant (Fig. 6D). A similar comparison of local tumor growth at the sites of injection revealed essentially no difference between the two groups (data not shown).

Histological examination of lung metastases that developed within 14-21 days following fat-pad injection of 4T1-shCon and 4T1-shEcad cells, demonstrated that loss of E-cadherin from the knocked-down cells was retained in the metastases, which were more scattered in the lung, compared to the 4T1-shCon cells, which showed a tendency to develop next to blood vessels (Supplementary Fig. S7).

To determine whether the higher metastatic activity of the WT 4T1 cells could be attributed to the more efficient dissemination of these cells from the primary tumor, or to their more effective extravasation and infiltration into the lung tissue, we injected either one or the other cell type intravenously into two groups of mice (Fig. 7A), and monitored lung bioluminescence using an IVIS imager. While both cell types initially accumulated in the lungs at similar rates (Fig. 7A, Days 0 and 3), by 8 days post-injection, the level of lung metastases formed by the 4T1-shCon cells was over 3.5-fold higher, compared to that of the 4T1-shEcad cells (Fig. 7A, Day 8, and B). Statistical analysis of these data suggested that the difference between the groups is highly significant (Fig. 7B). These results imply that the tether-forming 4T1 cells infiltrate more efficiently from the blood to the lungs, compared to the E-cadherin knockdown, tether-deficient cells.

**Figure 7:**
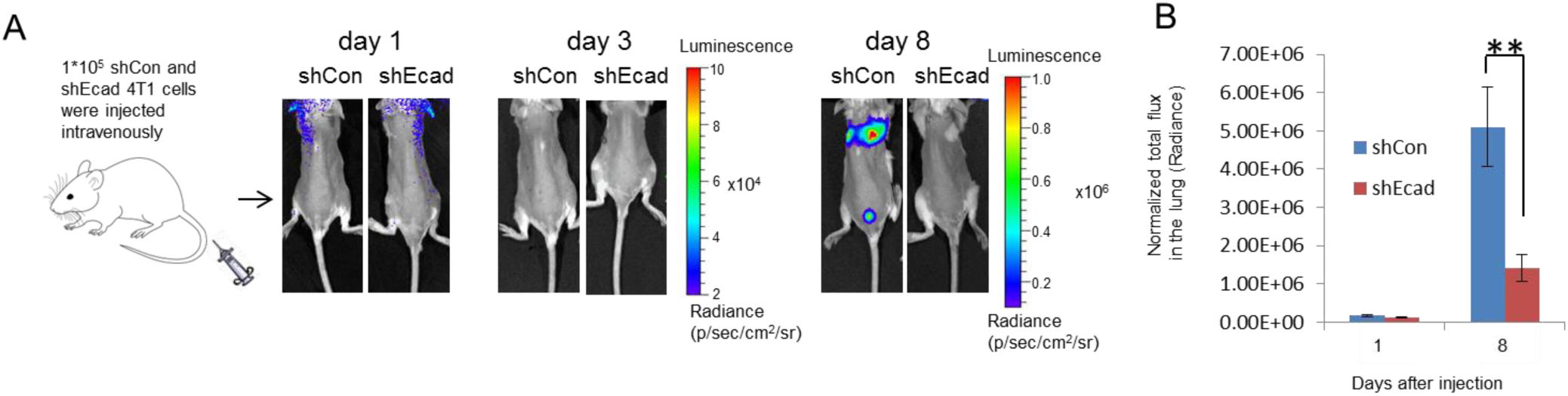
Knockdown of E-cadherin significantly reduces lung metastases following intravenous injection of 4T1 cells into BALB/c mice. (A) BLI imaging at 1, 3, or 8 days post-injection of 1*10^5^ 4T1-luciferase shCon or shEcad cells into the tail vein of BALB/c mice. (B) Quantification of total flux in the lungs of mice injected with 4T1-luciferase shCon or shEcad cells. Histograms indicate a major reduction in the development of lung metastases, in the E-cadherin knockded-down cells. Statistically significance of the differences between the total flux in the lungs of the mice, injected with the knocked-down and control cells was determined using two-way ANOVA, based on three independent experiments, n=11 in each group (**p= 0.001).

### Ex vivo imaging demonstrates the differential infiltration of 4T1 control and -shEcad cells into a monolayer of stromal fibroblasts

To directly compare the extent of invasiveness of 4T1 WT to that of the E-cadherin-deficient 4T1 cells, we developed an *ex vivo*, live-cell infiltration assay, whereby control and E-cadherin knockdown cells tagged with GFP, 4T1-GFP (Con) or 4T1-shEcad-GFP (shEcad), respectively, and mouse embryonic fibroblasts expressing red fluorescent protein (MEF-RFP) were cultured in parallel compartments of a silicone device attached to a culture plate (for details, see Experimental Procedures). The choice in MEF cells was based on previous experience demonstrating a differential susceptibility to penetration by invasive carcinoma cells. When the cells in both compartments reached confluence, the silicone inserts were removed, enabling the cancer and stromal cells to migrate towards each other.

As shown in Figure 8 and Supplementary Movie S10, during the first 7 hours post-insert removal, the Con and shEcad cells migrated into the cell-free space. Calculating the relative rates of “wound closure” by the two cell populations indicated that the average migration velocity of the shEcad cells was over 50% higher than that of Con (50.7 μm/hr, compared to 33.1 μm/hr, respectively; Fig. 8A and B t=4hr). Similar differences in migration velocity were also noted in the absence of stromal cells, or by tracking Con and shEcad cells in sparse cultures (data not shown). When the migrating cells (both Con and shEcad) reached the MEFs (at around t=12h-14h), the velocity of forward migration drastically dropped (Fig. 8B), reaching similar, low velocities in both cell types. Longer incubation (t=14h-38h) resulted in both cancer cell populations infiltrating into the “stromal domain”; yet the average rate of infiltration into the stromal monolayer by Con cells was 1.35-fold higher than that of shEcad (9.6 μm/hr compared to 7.1 μm/hr, respectively). Upon longer incubation infiltration (at t=38h-66h), velocities were increased further, the infiltration velocity of the Con cells was 16.0 μm/hr, while the shEcad cells’ mean velocity remained rather low (8.6 μm/hr). These results indicate that the E-cadherin expressing, collectively migrating 4T1 cells are more effective in infiltrating the stromal cell domain than the E-cadherin knockdown cells.

**Figure 8:**
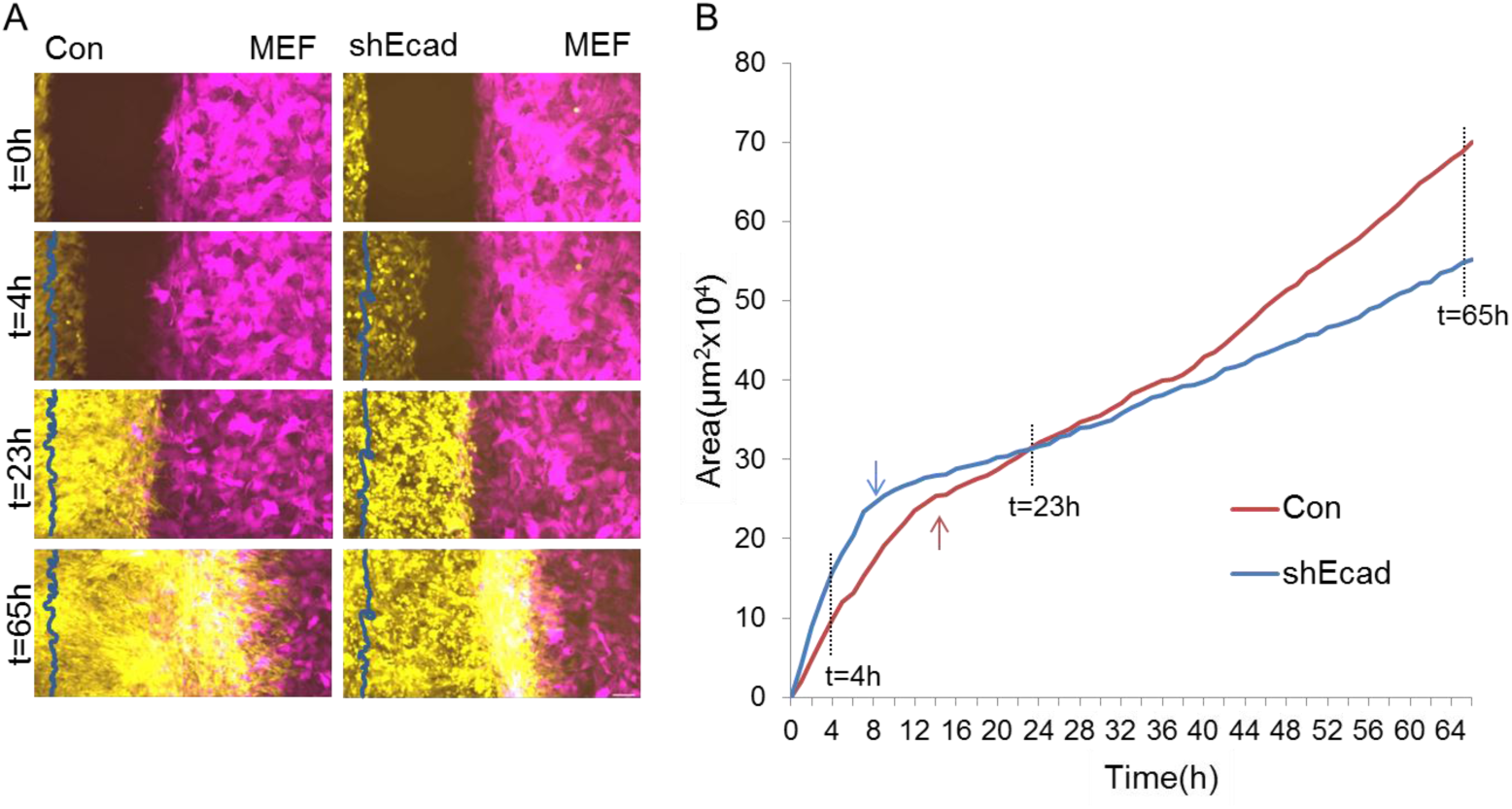
E-cadherin facilitates infiltration of 4T1 cell into stromal cells, *ex vivo*. GFP-expressing 4T1 cells (Con) or shEcad 4T1 cells (shEcad), and RFP expressing mouse embryo fibroblasts (MEF) were plated in two parallel compartments of a silicon insert (Ibidi^®^). When the cells within the two compartments reached confluence, the insert was removed, and the area between the two cell monolayers was imaged by live-cell video microscopy. (A) Snapshots taken at the indicated time points, showing the Con (left column, yellow) and shEcad (right column, yellow) and the MEF cells (magenta-colored). The vertical wavy line marks the position of the cancer cells at t=0. Scale bar: 100 μm. (B) A plot showing the positions of the “invasive front” of the shEcad (blue) and Con (red) over a period of 65 hours. The times at which the shEcad and Con 4T1 cells physically reach the fibroblasts are indicated by the blue and red arrows, respectively. Note that the initial migration rate of the shEcad cells is considerably faster than that of the control cells, while their infiltration into the stromal monolayer is considerably slower than that of the E-cadherin-expressing and tether-forming Con cells. For the full movie, see Supplementary Movie S10.

## Discussion

In this article, we addressed the involvement of a non-canonical form of collective migration, whereby invading cancer cells move, while being interconnected by a network of E-cadherin-mediated membrane tethers. We show here that these cell clusters display a highly invasive phenotype, despite the fact that they maintain their intercellular coherence. Most strikingly, we show that the loss of collective migration following suppression of E-cadherin expression in these cells leads to an increase in the rate of single-cell migration; yet it reduces cell invasiveness, both *in vivo* and *ex vivo*.

The interplay between collective- and single-cell migration in regulating cancer invasion and metastasis has attracted considerable interest in recent years ^2–4, 6^. The prevailing notion is that collective migration, which is primarily associated with local invasion into the stroma surrounding the tumor, requires protrusive forces produced by large cell collectives ^33^, and matrix degradation by secreted and invadopodia-associated matrix metalloproteinases ^34-36^. The development of cancer cell metastases, and their dissemination to distant organs, requires an additional step of cell individualization, which is primarily achieved by down-regulation of intercellular adhesions. Thus, cells are able to intravasate, undergo long-range translocation via blood vessels and lymphatics, and eventually extravasate and penetrate into target tissues. The small size of single cells, their deformability, and their coherent migratory persistence appear to drive their dissemination throughout the body ^17, 18, 21^.

The key process underlying cell individualization is EMT, whereby cell-cell adhesion molecules (e.g., E-cadherin) are down-regulated, and the cells express a series of classical mesenchymal markers ^7–9^. A common form of EMT involves the transient acquisition of a mesenchymal phenotype, induced by environmental factors secreted mostly by stromal and inflammatory cells. Evidently, when metastasizing cells reach an appropriate environment (e.g., the target organ), they may undergo mesenchymal-epithelial transition (MET) and reacquire epithelial characteristics, similar to those of the original tumor ^12, 13, 37^.

The findings presented in this article suggest that the invasive migratory phenotype displayed by 4T1 cells is regulated by a unique “EMT plasticity” process, whereby E-cadherin levels are moderately down-regulated to levels that enable, or even promote, loss of epithelial coherence, yet are high enough to support a special, “tether-mediated” mode of collective migration, that enhances stromal infiltration capacity *ex vivo*, and the efficient formation of lung metastases *in vivo*. Unlike classical EMT, induced by stromal and inflammatory cytokines ^38–40^, which commonly leads to massive loss of cell-cell adhesions, and acquisition of a mesenchymal migratory phenotype, mild reduction of E-cadherin levels leads to a mixed phenotype, with a blend of epithelial and mesenchymal properties (see also ^41–46^). The EMT plasticity displayed by 4T1 cells raises three main questions that deserve further discussion here; namely: (i) the mechanism underlying the reduction in E-cadherin levels in sparse cultures, or at the periphery of a tumor mass; (ii) the nature of the tethers, which support the collective migration of these cells; and (iii) the role of the residual E-cadherin in promoting invasion and metastasis, in culture models and *in vivo*.

The specific mechanisms underlying the metastatic properties of 4T1 cells have been debated for a long time. On the one hand, these cells display very aggressive metastatic properties; yet at the same time, they were reported to express high levels of E-cadherin and other epithelial markers, which are usually down-regulated in metastatic cancer cells ^30-32^. This apparent inconsistency is resolved by the results presented here, showing that the level of E-cadherin expressed in these cells is tightly regulated by cell density. The reduction of E-cadherin protein levels is modest, on the order of 50%; yet it has a major effect on cell migration, enabling a fast, tether-mediated collective and invasive migration, which is radically different from both the largely static cells in dense 4T1 cell islands, as well as from E-cadherin knockdown cells, which lose collectivity altogether, and display poor capacity to invade and metastasize.

We further demonstrate that suppression of E-cadherin levels in sparsely-plated cells occurs at the post-transcriptional level, as it is manifested at the protein level, without affecting the E-cadherin mRNA level. In this respect, the plasticity of the EMT-like state of sparse 4T1 cells differs from those cases of EMT induced by stromal or inflammatory factors, that commonly lead to a more substantial suppression of E-cadherin ^38, 47^, and exert their effects at the transcriptional level. Thus, epithelial cancer cells which express low levels, or no, E-cadherin (e.g. H1299 and MDA-MB-231), fail to form stable cell-cell tethers (see Fig. S1 and ref. ^48^).

The mechanism underlying the downregulation of E-cadherin protein levels in 4T1 cells is unclear; yet the phenomenon is consistent with the reduction in E-cadherin levels found in 4T1 cells cultured in on a 3D Matrigel matrix ^32^, and with the increased tendency of cancer cells located at the periphery of epithelial islands, to more readily undergo cytokine-induced EMT ^49^, suggesting that cadherin-mediated adhesion might suppress EMT. These results are also consistent with the capacity of 4T1 cells to form E-cadherin-dependent tethers, and migrate collectively in 3D collagen gels. It is noteworthy that a density-dependent, EMT-like response is not only manifested by 4T1 cells: a similar process was reported for a variety of other cell lines (e.g., SW480, colon cancer cells, and MCF10A breast cells), all of which undergo major phenotypic, EMT-like changes, including low levels of E-cadherin, when plated sparsely ^50, 51^. The capacity of moderate levels of E-cadherin to drive tether-dependent collective migration was also demonstrated here, by the controlled induction of E-cadherin in cells that lack E-cadherin expression (H1299), thereby switching their migratory behavior from “mesenchymal” to collective. Similarly, partial suppression of E-cadherin activity in tightly adherent epithelial cells (MCF10A), reduces the formation of tight cell-cell junctions and induces the development of tethers, and acquisition of fast collective migration.

The moderate reduction in E-cadherin levels in 4T1 cells leads to a non-canonical form of collective cell migration, mediated by thin membrane tethers. These tethers appear to be flexible, enabling cells to actively migrate, yet also robust enough to “restrain” the cells, and prevent loss of cluster coherence. Our initial morphological observations raised the possibility that the tethers are actually membrane nanotubes that interconnect the cytoplasms of neighboring cells ^52-54^. This possibility proved to be wrong by co-plating 4T1 cells expressing different fluorophores (RFP or GFP), and showing that there was no fluorophore transport from one cell to the other (data not shown). The robustness of the 4T1 tethers may be attributed to the fact that they contain microfilaments, microtubules and vimentin-based intermediate filaments (Fig. 3), much like the previously-described cytoskeletal tethers ^55^, which are quite mechanically stable ^56, 57^.

An intriguing question concerning the tethers’ role in maintaining migration collectivity is whether their ability to prevent cluster dispersal is merely passive, or whether they are capable of actively stimulating the departing cells to migrate back to their original locations. Careful examination of time-lapse movies (Fig. 2A and B) demonstrated that both the ”outward” and “inward” migrations were active processes, which proceeded at nearly identical speeds. We have no direct evidence of the mechanism underlying the tethers involvement in the “directional switch” of the lamellipodium, but previous studies have indicated that E-cadherin can regulate cell polarization and directional migration (e.g., ^58, 59^). Whether a similar mechanism is involved in regulating the internal cellular trafficking within the tether-interconnected clusters, remains unclear.

An intriguing feature of EMT plasticity ^42, 49, 60, 61^, particularly in 4T1 cells, is the requirement that E-cadherin support an invasive and metastatic phenotype. The prevailing paradigm concerning the role of “classical EMT” in cancer development is that the dramatic decline of cell adhesion, coupled with acquisition of mesenchymal migratory characteristics, enable cells at the tumor’s periphery to invade surrounding tissues. The results reported herein indicate that the mode and level of decline of E-cadherin can exert very different effects on the metastatic process. The initial reduction in E-cadherin levels due to low cell density indeed promotes invasive collective migration, as discussed above. On the other hand, further suppression of E-cadherin levels by its specific knockdown, which led to an overall 85% reduction of E-cadherin levels (compared to densely plated 4T1 cells), suppressed the development of lung metastases and cell infiltration into the stroma, despite the fact that their intrinsic migration rates on a fibronectin-coated substrate were considerably higher than that of control cells. This notion is in line with several recent reports ^30, 47, 62^, demonstrating that the majority of cancer cells still retain their epithelial morphology and migrate as clusters, throughout the metastatic process.

Taken together, these findings suggest that the residual E-cadherin expressed in sparsely plated 4T1 cells, though apparently insufficient for supporting robust adherens junction formation can, nonetheless, enable tether-mediated collective migration, and enhance the infiltration of the cancer cells into a stromal cell layer, and the formation of lung metastases. We still lack a definitive explanation as to the molecular underpinnings of this phenomenon, but would propose two possibilities, the first involving heterotypic interactions (e.g., E-cadherin/N-cadherin; see ^63, 64^)] between 4T1 cells and the surrounding fibroblasts; and the second, related to the potential contribution of the tethers to the infiltration process, enabling individual, tether-interconnected cells within a cluster to explore the ECM, identify migratory pathways, and migrate into them, along with their associated cells, thus enhancing metastatic infiltration. These potential mechanisms are currently being explored.

## Methods

### Antibodies, plasmids and reagents

Antibodies used in this study were obtained from the following sources: anti α-tubulin (T9026) and β-catenin (C2206), were purchased from Sigma-Aldrich (St. Louis, MO, USA); cingulin antibodies were kindly provided by Prof. Sandra Citi, University of Geneva, Switzerland; and anti E-cadherin (610181), was obtained from BD Transduction Laboratories (San Jose, CA, USA). Antibodies to vimentin (mouse, monoclonal) were locally produced. Staining of F-actin was performed using fluorescein isothiocyanate-labeled phalloidin (Sigma-Aldrich). Secondary antibodies used in this work included: goat anti-mouse IgG conjugated to Alexa Fluor 488 (Invitrogen™, Carlsbad, CA, USA), goat anti-mouse IgG conjugated to Cy5 (Jackson ImmunoResearch Laboratories, West Grove, PA, USA), and goat anti-rabbit IgG conjugated to cy3 (Jackson ImmunoResearch Laboratories).

Tetracyclin-inducible E-cadherin promotor plasmid was a kind gift of Prof. Barry M. Gumbiner, Seattle Children’s Research Institute, Seattle, WA, USA).

Lentivirus encoding GFP was kindly provided by Dr. Ravid Straussman (Weizmann Institute of Science). 4T1-luciferase plasmids: pLenti PGK V5-LUC Neo was obtained from Addgene (Cambridge, MA, USA). pLP1, pLP2, and pLP/VSVG were all obtained from Invitrogen™. E-cadherin knockdown: TRC cdh1 shRNA - TRCN0000042578 (GE Healthcare Dharmacon, Lafayette, CO, USA); pCMV-VSV-G (helper plasmid) and pHRCMV-8.2ΔR (packaging plasmid) were both kindly provided by Prof. Yardena Samuels (Weizmann Institute of Science). Mission pLKO.1-puro (Sigma-Aldrich) was used as negative control vector, to create 4T1-shCon cells. Other reagents: Bovine fibronectin was purchased from Biological Industries (Beit HaEmek, Israel); and EGTA, from Sigma-Aldrich. Cells were transfected using Lipofectamine2000^®^ (Invitrogen™), according to the manufacturer’s instructions.

### Immunofluorescence staining

For staining that included tubulin: Cells were fixed with 2.5% PFA, 0.2% glutaraldehyde in PBS containing 0.2% Triton X-100 for 20 min, washed 3 times in PBS, washed for 15 min with PBS and sodium borohydride, then washed again 3 times in PBS.

For staining that didn’t involve tubulin/vimentin, cells were permeabilized with 3% PFA in PBS containing 0.5% Triton X-100 for 2 min, then fixed with 3% PFA in PBS for 20-30 min, and washed 3 times in PBS.

For vimentin staining: cells were fixed with -20°C methanol for 10 min, washed 3 times with -20°C acetone, then washed 3 times with PBS.

### Fluorescence microscopy and live-cell imaging

Live-cell imaging and sample examination were performed using a DeltaVision Elite microscope system (GE Healthcare Bio-Sciences, Pittsburgh, PA, USA).

Fluorescence images were acquired at 60x magnification (60x/1.42NA objective), by a CoolSnap HQ2 CCD camera^®^ (Roper Scientific, Planegg, Germany). Time-lapse movies were acquired using phase contrast or interference contrast microscopy, with 10x/0.3NA objective, at time intervals of 1/5/15 min between frames, as indicated.

### Cell migration on 3D matrices

Collagen-based 3D matrix was prepared by mixing minimal essential Eagle’s medium (MEM) (Flow Laboratories, McLean, USA) and 19% rat tail collagen type 1 (BD 354249), at pH 7.5 (protocol was modified from ^65^). 4T1 cells were trypsinized, mixed with the collagen gel, which was placed on a 35mm glass-bottom dish (No. 0, Uncoated MatTek™, Ashland, MA, USA), and covered by a glass coverslip, that was fixed to the dish by a biocompatible glue (Picodent twinsil^®^,Wipperfürth, Germany). Fresh medium was added to the plate and live cell imaging was initiated ~3 hr thereafter, focusing on cells located at least 50 μm away from the top or bottom glass surfaces.

### High resolution scanning electron microscopy (UR-SEM) imaging

Cells were grown on a glass surfaces, coated with 10μg/ml fibronectin (FN), and fixed with 2.5% PFA and 2.5% glutaraldehyde in 0.1M cacodylate buffer, pH 7.2, followed by post fixation with 1% OsO4 in the cacodylate buffer. The fixed cells were dehydrated in increasing ethanol concentrations (30%, 50%, 70%, 96%, and 100%) in Milli-Q^®^ water, and critical-point dried using BAL-TEC CPD 030). Dry samples were coated with a thin layer of carbon in Edwards carbon coater and visualized using secondary electron (SE) and back-scattered electron (BSE) detectors in a high-resolution Ultra 55 SEM^®^ (Zeiss, Germany), using a 30 μm aperture size, a working distance of 5.0 mm, and 3.0 kV voltage.

### Cell Cultures

Cells used in this study included: 4T1 (Murine mammary carcinoma cell line, ATCC^®^ CRL-2539™), MDCK (Canine normal kidney epithelial cell-line, ATCC^®^ CCL-34™), H1299 (human non-small cell lung carcinoma cell-line, ATCC^®^ CRL-5803™), and MCF10A (human non-tumorigenic breast epithelial cell line, ATCC^®^ CRL-10317™). The 4T1, MDCK and H1299 cells were cultured in PRMI1640 (Gibco™, Waltham, MA, USA) plus 10% FBS, 2mM L-glutamine (Biological Industries), and 1% Pen-Strep (Biological Industries). MCF10A cells were cultured in DMEM/F-12 (Biological Industries), plus 5% horse serum (Gibco™), 0.5 μg/ml hydrocortisone (Merck Millipore, Billerica, MA, USA), 0.1 μg/ml Cholera toxin (Sigma-Aldrich), 10 μg/ml insulin (Biological Industries), 1% Pen-Strep, and 10 ng/ml EGF (Sigma-Aldrich).

### Animal studies

Analyses of primary tumor growth and lung metastases were performed in 10-12 week-old female BALB/c mice. One-day pre-injection, mice were anesthetized using 2 mg/10 gr body weight sedaxylan, 10 mg/10 gr body weight Ketamine, and shaved using a shaving machine and Orna 19 (hair removal cream, Alpha Cosmetics, Israel).

Primary tumor injection: 2.5 *10^4^ 4T1-luciferase (shCon or shEcad) cells in 50 μl PBS were injected into the mammary fat pad.

Intravenous injection: 1*10^5^ 4T1-luciferase (shCon or shEcad) cells in 150 μl PBS were injected into the tail vein. For visualizing tethers in lung metastases, 1*10^6^ 4T1-GFP cells were injected intravenously, animals were sacrificed, and a small metastatic nodules in the lungs were immediately imaged, using 2-photon microscopy (LSM 880, Zeiss; 20x objective).

Bioluminescence imaging (BLI): Prior to mice imaging, 50 μl of 30 mg/ml luciferin (Regis Technologies, Inc., Morton Grove, IL, USA) was injected intraperitoneally. After 7 minutes, mice were anesthetized using isoflurane (AbbVie, Inc., North Chicago, IL, USA), delivered with oxygen from a precision vaporizer. Non-invasive BLI was performed using a Xenogen IVIS Spectrum (PerkinElmer, Inc., Waltham, MA, USA).

### Statistical Analysis

Quantitative data for statistical analysis were expressed as mean ± SEM (shown as error bar) from at least two independent experiments. The significance of the difference between the metastatic activity of shCon and shEcad 4T1 cells (Fig 5, G and H) was calculated using the Student’s t-test. Significance between the different mice treatment groups was analyzed with two-way ANOVA on two (fat pad) or three (IV; tail vein) separate experiments.

### Quantitative real-time PCR (QRT–PCR)

Total RNA was isolated using an RNeasy Mini Kit (Qiagen, Hilden, Germany), according to the manufacturer’s protocol. A 2 μg aliquot of total RNA was reverse-transcribed, using a high-capacity cDNA reverse transcription kit (Applied Biosystems, Foster City, CA, USA). Quantitative real-time PCR (QRT–PCR) was performed with a OneStep instrument (Applied Biosystems), using Fast SYBR^®^ Green Master Mix (Applied Biosystems). Gene values were normalized to a HPRT1 or GUSB housekeeping gene. The following primers were used: E-cadherin: F 5′ AGCTCTAAGGACAGTGGTCAT 3′, R 5′ CAGTGCTTTACATTCCTCGGT 3′. HPRT1: F 5′TCCATTCCTATGACTTAGATTTTATCAG 3′, R 5′ AACTTTTATGTCCCCCGTTGACT 3′. GUSB: F 5’ CCGACCTCTCGAACAACCG 3’ R 5′ GCTTCCCGTTCATACCACACC 3’.

### Immunoblotting

Whole-cell lysates were prepared using RIPA buffer (150 mM NaCl, 1mM EDTA, 1% Triton X-100, 0.1% sodium deoxycholate, 0.1% SDS, 50 mM Tris, pH 8.0, containing 1mM PMSF) (Sigma-Aldrich). Protein concentrations were determined using the Pierce^®^ BCA protein assay kit (Thermo Fisher Scientific, Waltham, MA, USA), and samples were resolved on 10% SDS-Page gels. Blots were probed with antibodies to E-cadherin and α-tubulin (control) 1:2000, and developed using SuperSignal^®^ ECL reagents (Thermo Fisher Scientific). Lanes were extracted from the same gel.

### Infiltration cell-tracking assay

Tissue culture dishes were pre-coated with 10 μg/ml FN. 4T1-GFP (Con)/ shEcad, and MEF-RFP cells (3 *10^4^) were grown in the two wells of a silicon insert (Ibidi^®^, Germany), until reaching confluence. The insert was then removed, and the area between the wells was subjected to live cell imaging. The migratory front of the cancer cells was marked, manually and the migration rates were calculated using ImageJ software (National Institutes of Health, Bethesda, MD, USA).

## ACKNOWLEDGEMENTS

We would like to acknowledge support by the European Union Seventh Framework Program ERC Advanced Grant under grant agreement no. 294852-SynAd, and by the Israel Science Foundation, grant no. No. 3001/13 (both to BG). The authors are grateful to Barbara Morgenstern for her expert help in the style editing of this manuscript. We thank our colleagues at the Weizmann Institute of Science for their helpful guidance, including: Dr. Elena Kartvelishvily (Department of Chemical Research Support), for her assistance with the scanning electron microscopy imaging; Prof. Zvi Kam (Department of Molecular Cell Biology) for developing the “nearest neighbor” algorithm; and Dr. Ron Rotkopf (Biological Services Unit) for assisting with statistical analysis of the mouse data. BG is the incumbent of the Erwin Neter Professorial Chair in Cell and Tumor Biology.

## Supplementary information

### Diversity of collective migration modes

To verify which cell lines migrate collectively, we used the “nearest neighbor” methodology, in which we seed cells sparsely, grow them to about 20% confluence, then fix them and stain their nuclei with DAPI. In each cell line, we measured the average distance between each cell nucleus, and its three closest neighbors (Fig. S1B). By plotting the percentage of cells against the average three “nearest neighbors” distances, we noticed that 4T1, MCF10A, and MDCK cells create sharp peaks between 20-50 μm. This finding indicates that these three cell lines create cell clusters in which the distance of one cell from its neighbors is small, while H1299 cells, which migrate in a non-collective manner, were distributed quite homogenously. Notably, less than 2% of 4T1 cells contain more than one nucleus, supporting our contention that the cells migrate collectively. Furthermore, the low cell density in the plate indicates that cell clusters were not created by being plated densely. We also confirmed that the cells were thoroughly suspended and seeded as single cells to minimize initial cell-cell interactions.

## SUPPLEMENTARY FIGURE LEGENDS

**Figure S1:**
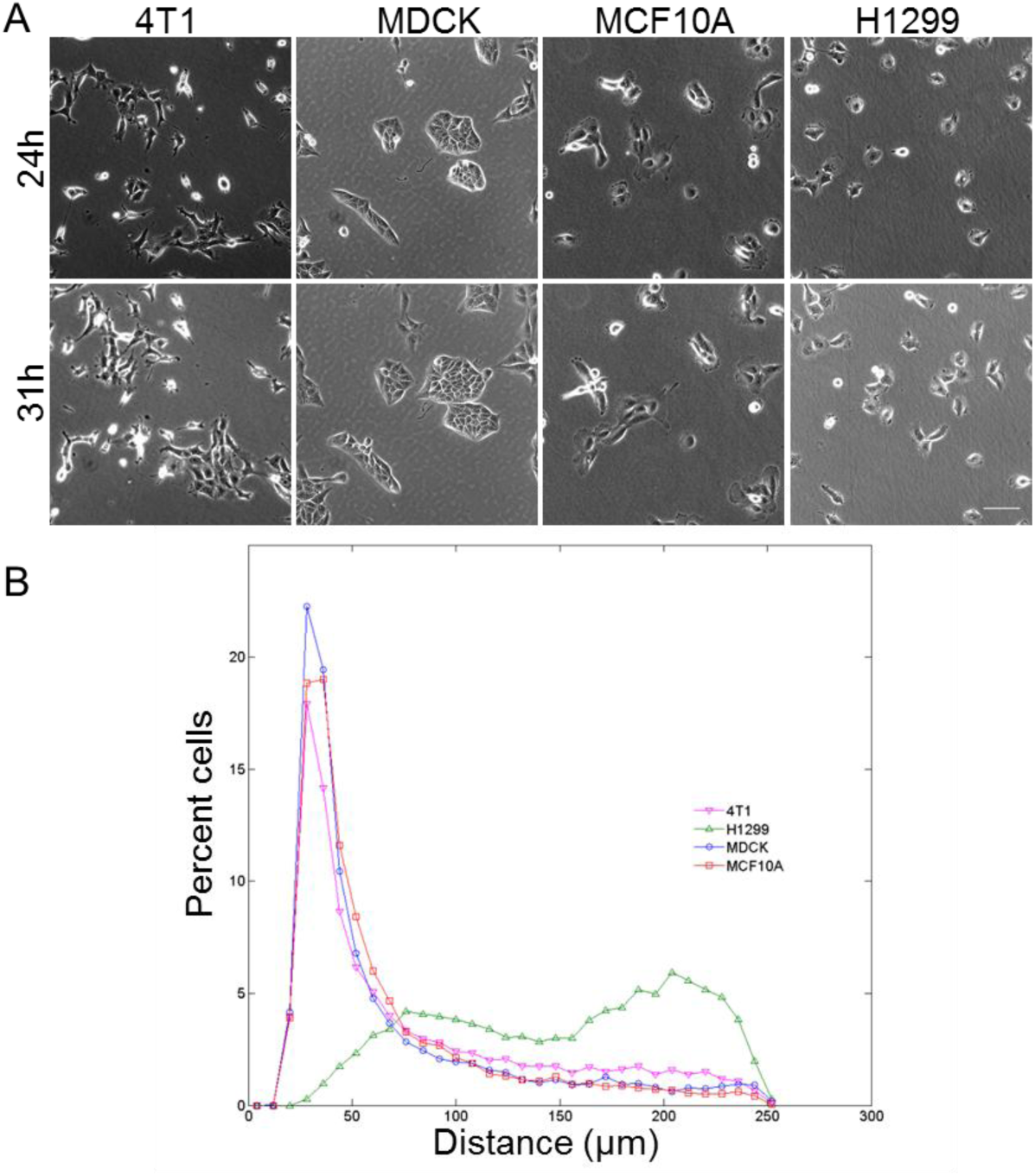
Scoring of collective cell migration displayed by different epithelial cell lines, using a “nearest neighbor” collectivity scoring. (A) Different epithelial cell lines (4T1, H1299, MDCK, MCF10A) were sparsely plated on tissue culture dishes, coated with 10 μg/ml fibronectin (FN) and imaged, using phase-contrast microscopy, 24 hrs and 31 hrs thereafter. Scale bar: 100 μm. (B) For “nearest neighbor” collectivity scoring, the four cell types shown in Panel A, were sparsely plated, and incubated for 24-48 hrs, after which they were fixed and stained with DAPI. The positions of nuclei were automatically recorded, and the average distance between each nucleus and its 3 nearest neighbors calculated. The plot indicates that MCF10A, MDCK and 4T1 maintain sharp peaks, within the range of roughly 20-30 μm between them, while H1299 cells, plated at an identical density, displays a broad peak, ranging from 50-240 μm, characteristic of single-cell migration. Notably, the collective translocation of epithelial islands varies greatly, as can be seen in Supplementary Movie S1.

**Figure S2:**
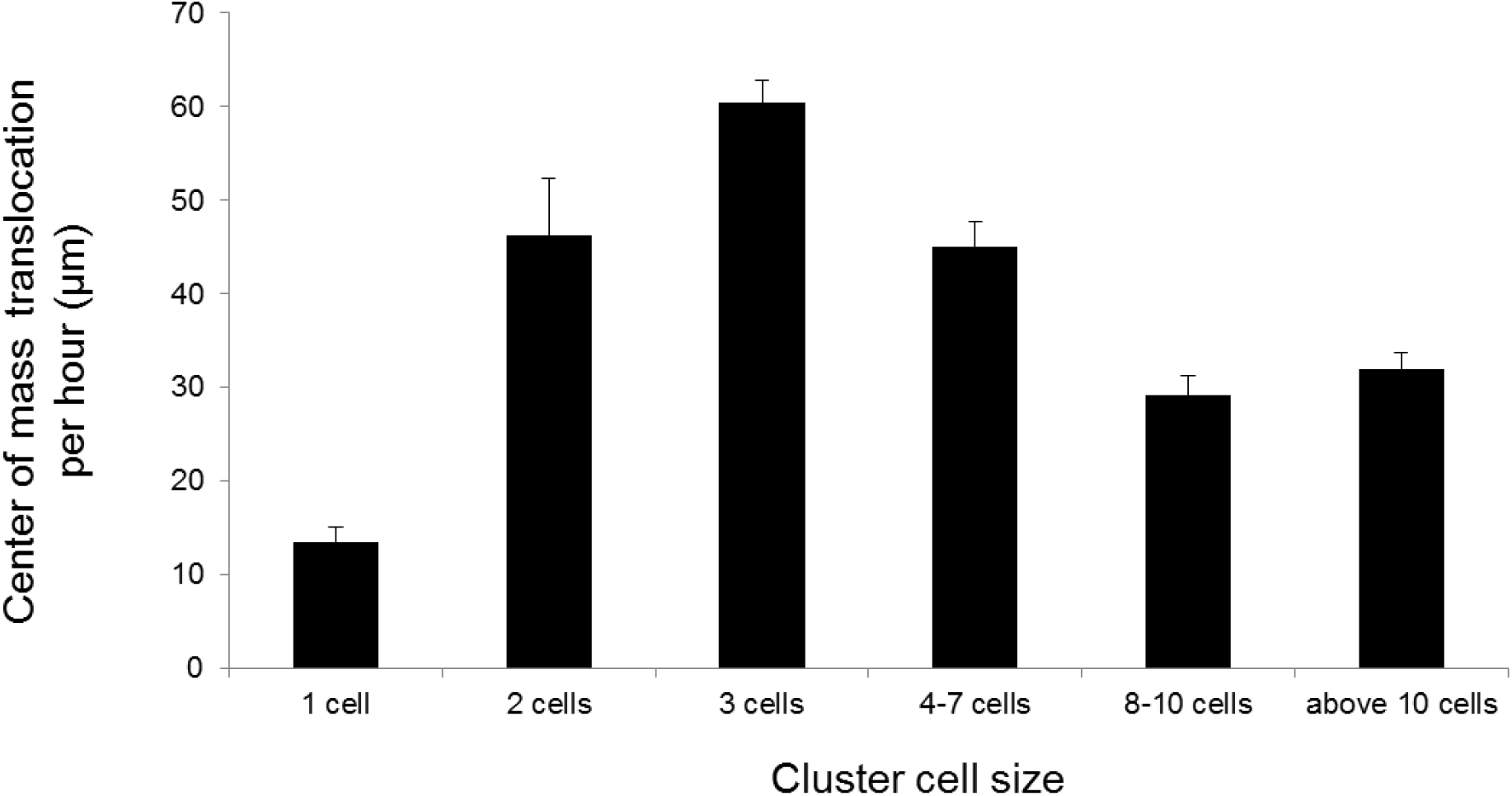
Small 4T1 cells clusters translocate faster, compared to large clusters or single 4T1 cells. 4T1 cells were cultured for 24 hrs in a tissue culture dishes, coated with 10 μg/ml fibronectin (FN). Cell migration was then monitored by live-cell imaging (5 min interval between frames). In these movies, we manually measured center-of-mass translocation of different-sized cell clusters over time, using ImageJ software. Each histogram represents the tracking of at least three different-sized cell clusters, in two independent experiments.

**Figure S3:**
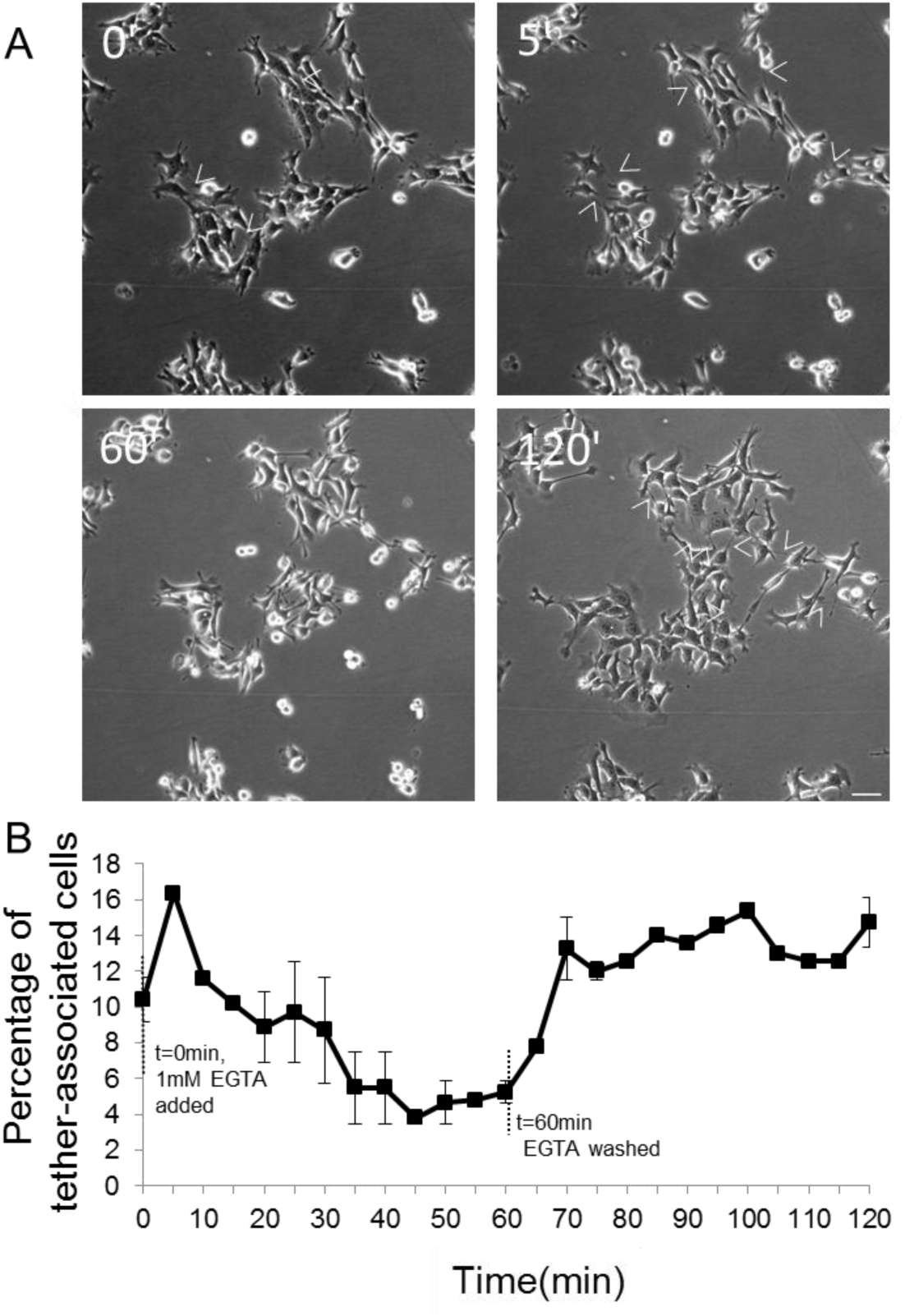
4T1 inter-cellular tethers are calcium-dependent. 4T1 cells were cultured for 24 hrs in tissue culture dishes, coated with 10 μg/ml fibronectin (FN). Cells were imaged just before (t=0), or at different time points after the addition of 1mM EGTA to the medium. (A) EGTA treatment induced cell contraction, leading to the formation of multiple short-lived tethers (t=5 min), which essentially disappeared upon further incubation (t=60min). Replacement of the medium at t=60 min with normal, (Ca^++^-containing) medium led to gradual increase in tether number, reaching normal levels by 60-120 min. Arrowheads in panel “5 min” and “120 min”, point to ruptured tether and newly-formed tether, respectively. Scale bar: 50 μm. (B) Percentage of tether-associated cells was measured at various time points during calcium chelator incubation (0-60 min) and after calcium restoration (60-120 min).

**Figure S4:**
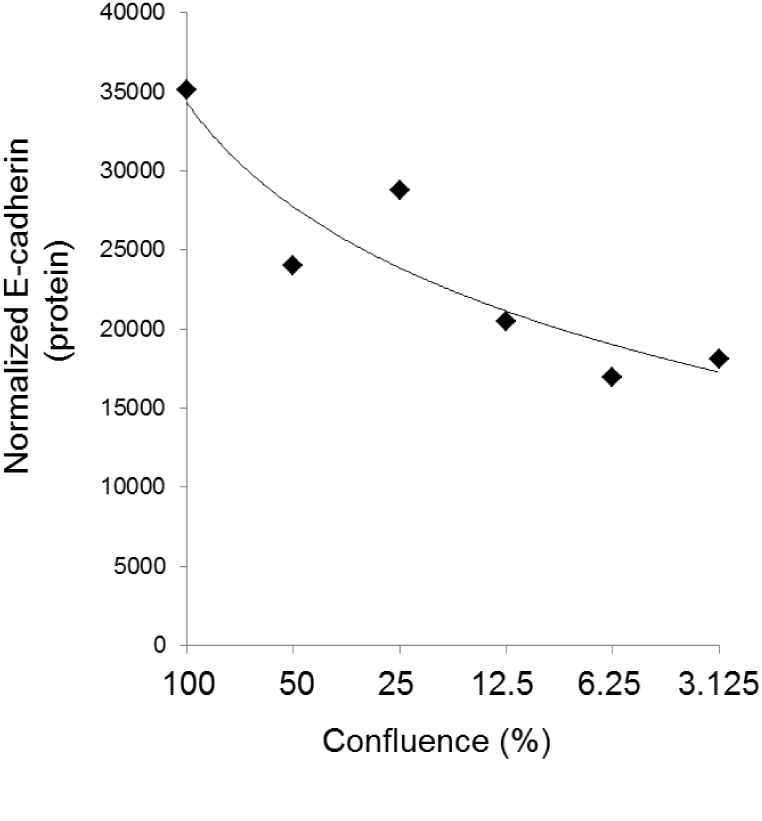
E-cadherin protein levels are affected by plating density of 4T1 cells. Serial dilution of plated 4T1 cells/cm^2^ from 8.9*10^4^ (100% confluence) to 2.8*10^3^ (3.125% confluence) followed by 24 hr incubation, affected E-cadherin levels, as measured by Western blot analysis. Notice that at ~10% confluence or lower, E-cadherin levels display a twofold lower E-cadherin protein levels, compared to those of cells plated at 100% confluence.

**Figure S5:**
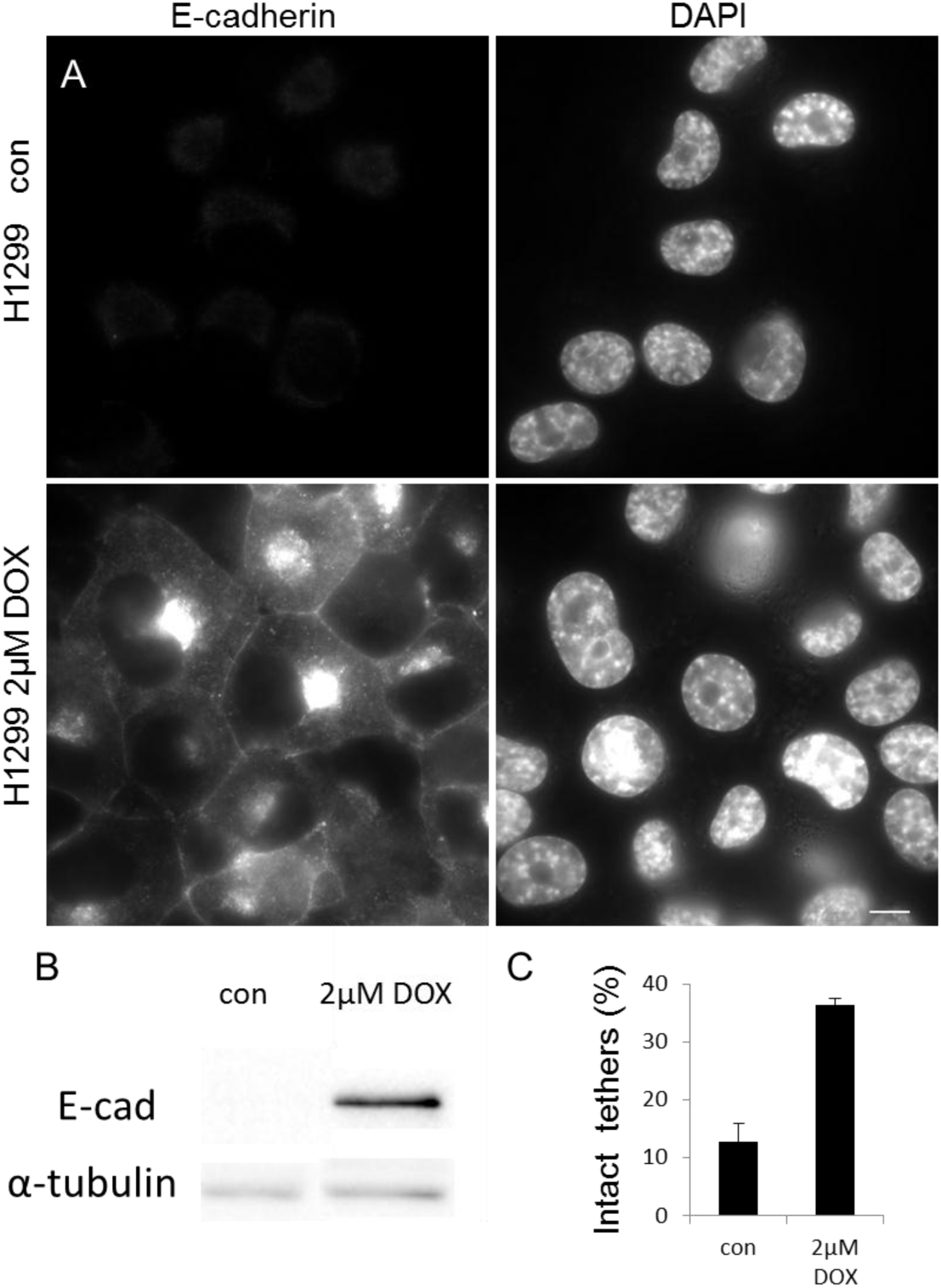
Over-expression of E-cadherin in H1299 cells induces the formation of stable tethers in these cells. H1299 cells expressing a tetracycline-inducible E-cadherin promotor, were cultured for 24 hrs in tissue culture dish with/without doxycycline (DOX, 2μM). (A) E-cadherin expression was visualized by immunolabeling (A) or Western immunoblotting (B). Notice the increased expression of E cadherin following DOX induction (A and B), and the major increase in tether formation (~2.8 fold) shown in C.

**Figure S6:**
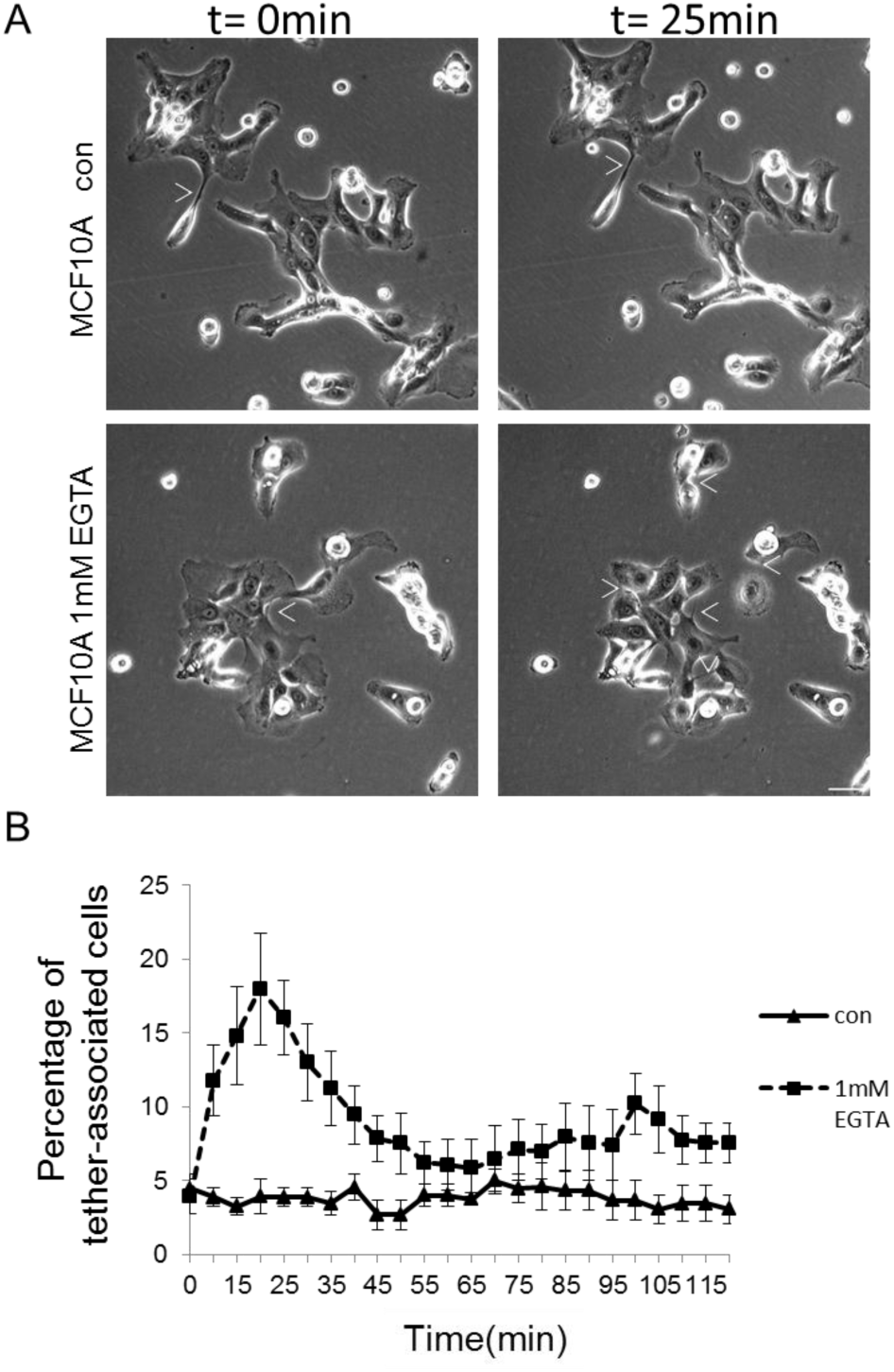
Reduction of calcium levels in the medium, leads to tethers formation by MCF10A cells. MCF10A cells, which, normally, form robust adherens-type junctions (not mediated via tethers). were plated for 24 hrs in a tissue culture dish coated with 10 μg/ml fibronectin (FN), after which 1mM EGTA was added to the medium (t=0). (A) EGTA treatment induced cell contraction and formation of many short-lived tethers (t=25 min), which essentially disappeared upon further incubation. Arrowheads point to the tethers detected at the indicated timepoints. Scale bar: 50 μm. (B) Percentage of tether-associated cells, measured at various timepoints during EGTA treatment (0-120 min).

**Figure S7:**
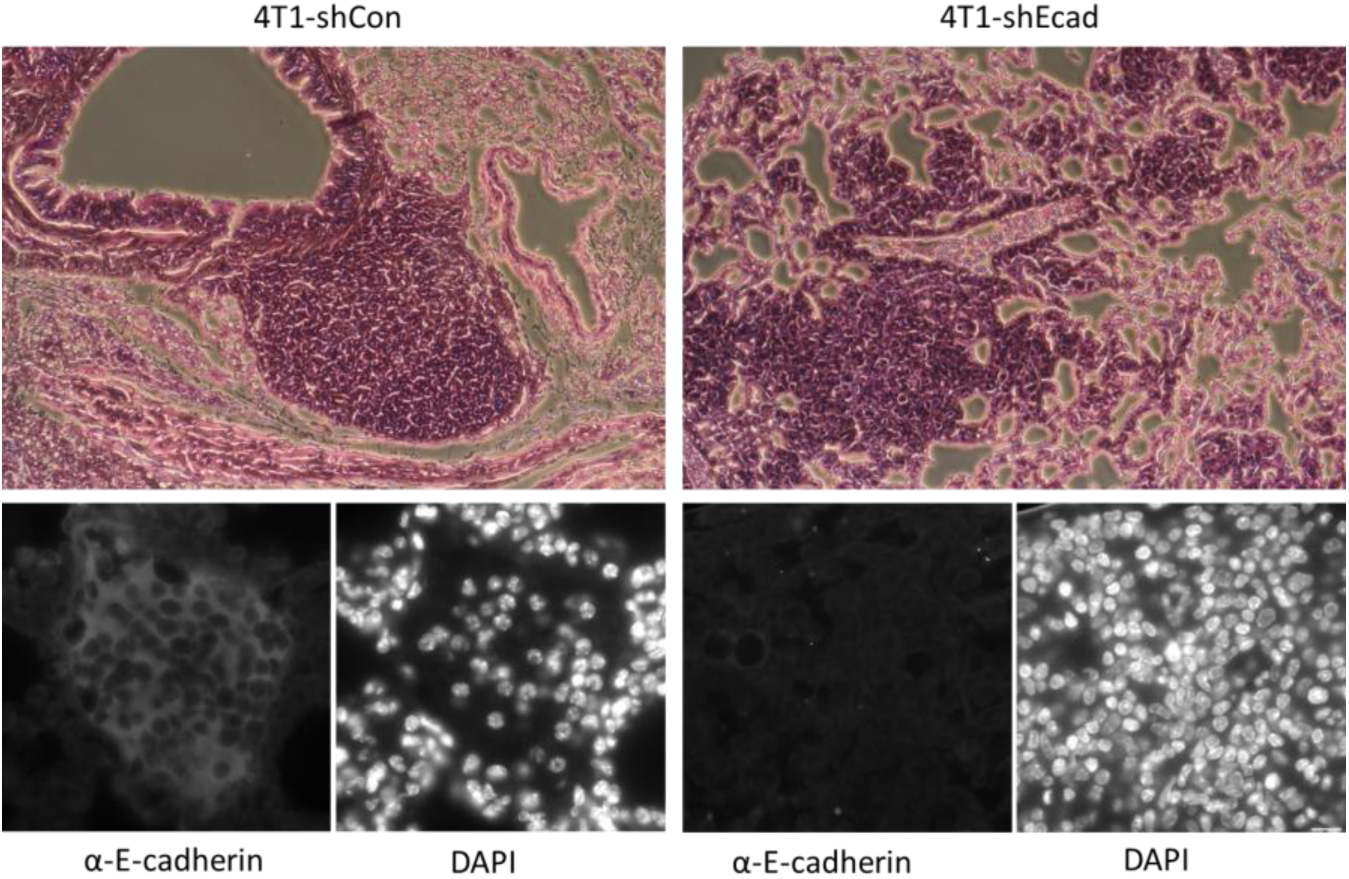
Lack of E-cadherin expression persists in lung metastases following injection of 4T1-shEcad cells into the mammary fat pads of BALB/c mice. Representative histological sections of metastases-containing lungs, examined 14-21 days after fat-pad injection of 4T1-shCon and 4T1-shEcad cells. Immunolabeling demonstrated that while E-cadherin staining is clearly visible in 4T1-shCon cells, no E-cadherin staining is found in 4T1-shEcad mice lung sections. Notice that 4T1-shCon metastasis tended to grow in clusters often located in close proximity to blood vessels, primarily at the lung’s periphery, while 4T1-shEcad cells tended to grow in a more disorganized manner throughout the lung.

## SUPPLEMENTARY MOVIE LEGENDS

### Supplementary Movie S1

Different epithelial cell lines (4T1, H1299, MDCK, MCF10A) were sparsely plated on tissue culture dishes coated with 10 μg/ml fibronectin (FN). Twenty-four hrs post-seeding, cells were imaged, using live cell vide microscopy (with phase-contrast optics; 10x/0.3NA objective; time interval between frames= 5-min) Note the collective mode of migration of MDCK (slow), MCF10A (medium) and 4T1 (fast), and the tendency of H1299 cells to migrate individually. Scale bar: 10 μm.

### Supplementary Movie S2

4T1 cells were seeded sparsely on a tissue culture dish, coated with 10 μg/ml fibronectin (FN), and immediately subjected to live-cell imaging,. Cells were imaged every 5 min, using a 10x 0.3NA phase-contrast objective. Scale bar: 10 μm.

### Supplementary Movie S3

4T1 cells were seeded on a tissue culture dish coated with 10 μg/ml fibronectin (FN) for 24 hrs, after which the cells were imaged every 5 min for a total of 2.75 hrs, using phase-contrast microscopy with a 10X0.3NA objective. Scale bar: 10 μm.

### Supplementary Movie S4

4T1 cells were cultured for 24 hrs on a tissue culture dish coated with 10 μg/ml fibronectin (FN), after which 1mM EGTA was added to the medium. Arrows point to the tethers tracked during the movie, both prior to and immediately after addition of EGTA. After 60 mins, the medium was replaced by fresh, calcium-containing medium and cells were further imaged for 60 mins with 1 min interval between frames. Live cells imaging was conducted, using phase-contrast microscopy with a 10x 0.3NA objective. Scale bar: 50 μm.

### Supplementary Movie S5

H1299 cells expressing a tetracycline-inducible E-cadherin were cultured for 24 hrs in a tissue culture dish with (DOX) or without (H1299 con) 2μM doxycycline. Cells were imaged every 5 minutes for a total of 495 mins, using phase-contrast microscopy with a 10x 0.3NA objective. Scale bar: 50 μm.

### Supplementary Movie S6

MCF10A cells were cultured for 24 hrs on a tissue culture dish coated with 10 μg/ml fibronectin (FN). Cells were then subjected to live cell video imaging, and 1mM EGTA was immediately added to the medium (MCF10A, 1mM EGTA). As controls, MCF10A cells in another plate, were tracked without the addition of EGTA (MCF10A con). Cells, on both plates, were imaged every 5 min for a total of 120 mins, using phase-contrast microscopy with a 10x 0.3NA objective. Scale bar: 50 μm.

### Supplementary Movie S7

4T1 shCon and -shEcad cells were seeded, separately, on tissue culture dishes coated with 10 μg/ml fibronectin (FN). Twenty-four hrs post-seeding, cells were imaged every 5 min for 8.25 hrs total, using phase-contrast microscopy with a 10X0.3NA objective. Note the tethers formed by 4T1-shCon cells, which connects them to neighboring cells, and the lack of tether formation from 4T1-shEcad cells. Scale bar: 10 μm.

### Supplementary Movie S8

4T1-shCon and 4T1-shEcad cells were cultured in 3D collagen gels, as described in Methods. After 3 hrs, the cells were tracked by microscopy, every 5 min, for a total of 3 hrs, using phase-contrast video microscopy, using a 10x/0.3NA objective. Scale bar: 50 μm.

### Supplementary Movie S9

4T1-GFP-shCon cells (1*10^6^) were intravenously injected into BALB/c mice. Seven days later, the animals were sacrificed, and their lungs were imaged using 2-photon microscopy (using 20x objective). 3D-reconstruction was performed using Imaris Bitplane^®^ 7.2 Software, clearly showing the presence of tethers, interconnecting neighboring cells. Scale bar: 10 μm.

### Supplementary Movie S10

GFP-expressing 4T1 cells (Con) or shEcad 4T1 cells (shEcad), and RFP expressing mouse embryo fibroblasts (MEF) were plated in two parallel compartments of a silicon insert (Ibidi^®^). When the cells within the two compartments reached confluence, the insert was removed, and the area between the two cell monolayers was imaged by live-cell video microscopy with 15 min intervals between frames, for a total of 66.75 hrs, using a 10X0.3NA objective. Scale bar: 10 μm.

